# Epigenetic trajectory predicts development of clinical rheumatoid arthritis in anti-citrullinated protein antibody positive individuals: Targeting Immune Responses for Prevention of Rheumatoid Arthritis (TIP-RA)

**DOI:** 10.1101/2024.10.15.618490

**Authors:** E. Barton Prideaux, David L. Boyle, Eunice Choi, Jane H. Buckner, William H. Robinson, V. Michael Holers, Kevin D. Deane, Gary S. Firestein, Wei Wang

## Abstract

The presence of anti-citrullinated protein antibodies (ACPAs) in the absence of clinically-apparent inflammatory arthritis (IA) identifies individuals “at-risk” for developing future clinical rheumatoid arthritis (RA). However, it is unclear why some ACPA+ individuals convert to clinical RA while others do not. We explored the possibility that epigenetic remodeling is part of the trajectory from an at-risk state to clinical disease. Cross-sectional differential methylation analysis at baseline revealed DMLs that distinguish the Pre-RA methylome from ACPA+ Non-converters. Genes overlapping these DMLs correspond to aberrant NOTCH signaling and DNA repair pathways in B cells. Longitudinal analysis showed that ACPA-Control and ACPA+ Non-converter methylomes are relatively constant. In contrast, the Pre-RA methylome remodeled along a dynamic “RA methylome trajectory” characterized by epigenetic changes in active regulatory elements. Machine learning revealed individual loci predictive of RA conversion. DNA methylation is a dynamic process in ACPA+ individuals at-risk for developing RA that later transition to clinical disease. In contrast, non-converters and controls have stable methylomes. The accumulation of epigenetic marks over time prior to conversion to clinical RA conforms to pathways that are associated with immunity and can be used to identify potential pathogenic pathways for therapeutic targeting and/or use as prognostic biomarkers.

## Introduction

Rheumatoid arthritis (RA) development is a process that begins years before the onset of clinically apparent signs and symptoms consistent with classifiable disease^1,2^. Current concepts suggest that mucosal immune responses, especially in the airway and gut^2,3^, lead to a break in self-tolerance. Subsequent development of anti-citrullinated protein antibodies (ACPAs) in mucosal sites such as the lung and then systematically are important steps in the transition from health to autoimmune disease^4^. ACPA detection in blood using anti-cyclic citrullinated peptide (CCP) antibody tests can identify individuals at high risk for developing RA, creating an opportunity to follow them longitudinally and understand pathogenic mechanisms^5^.

To study the immunologic events that lead to clinical RA, the Targeting Immune Responses for Prevention of Rheumatoid Arthritis (TIP-RA) Collaborative developed cohorts of ACPA positive (anti-CCP3+) and ACPA negative (anti-CCP3-) individuals and early RA patients^6^. Our initial cross-sectional study of peripheral blood cell populations identified immune dysregulation in at-risk individuals, including high ACPA production with multiple specificities, aberrant DNA methylation and antigen-specific responses to certain citrullinated peptides prior to onset of synovitis. Differences in DNA methylation between the anti-CCP3+ at-risk and early RA populations in that analysis were striking, suggesting that epigenetic marks might evolve over time. Previous studies in support of epigenetic evolution in clinical RA also showed that patients presenting with undifferentiated arthritis (UA) exhibit distinct DNA methylation profiles compared to healthy controls and that UA patients who later develop clinical RA could be distinguished from those who would not^7^.

To define the trajectory of DNA methylation abnormalities in at-risk individuals and determine at what point the epigenome becomes altered in an informative manner, our cross-sectional analysis was expanded, including longitudinal analyses to determine how the epigenome in peripheral blood immune cell populations is remodeled during the pre-RA period as individuals progress to clinical RA. The TIP-RA consortium now reports the results for anti-CCP3+ individuals at-risk for RA that have been followed for up to five years. We discovered that the methylome of individuals who will develop RA can be distinguished from anti-CCP3+ participants who do not convert to classifiable disease, which closely resemble anti-CCP3-individuals without IA involving critical pathways like NOTCH. Further, we discovered that the epigenomes in individuals remodel along a “rheumatoid arthritis methylome trajectory” over time, both prior to and including the time of first detection of clinical RA. We then developed and tested prediction models designed to identify the most important epigenetic loci associated with the conversion to clinical RA and that could lead to the development of prognostic tests and/or identify novel therapeutic targets.

## Results

### Study Design and Method Validation

Three groups were studied in TIP-RA cohorts: 172 anti-CCP3-“Controls” without IA, 97 anti-CCP3+ “At-Risk” participants and 62 anti-CCP3+ “Early RA” patients were enrolled in the TIP-RA cohort^6^. At-Risk and Control participants were followed for 5 years, with blood samples collected annually for each group, as well as at or immediately after the first identification of a swollen join consistent with synovitis (i.e. Clinical RA) (see reference 6 for a more complete description of the cohort and study). A subset of these individuals (Supplementary Fig. 1) were evaluated for DNA methylation in naïve and memory CD4+ T cells and B cells (69 Controls, 71 At-Risk and 29 Early RA). Over the course of the project, 21 anti-CCP3+ At-Risk participants developed clinical RA. These individuals are referred to as “pre-RA”, while the remaining 50 participants who did not develop RA are called “non-converters”.

There were no significant differences between At-Risk and Control participants with regard to age, sex, history of ever or current smoking, self-reported first-degree relative RA status, the presence of the HLA-DR shared epitope associated with RA, and level of high sensitivity C-reactive protein (hsCRP) (Table 1). However, the At-Risk individuals exhibited significantly higher prevalence of serum Immunoglobulin (Ig) M, IgG and IgA rheumatoid factor (RF) at baseline (*p* < 0.05, BH-adjusted Welch Two Sample t-test and Chi-Square test).

**Table 1.**
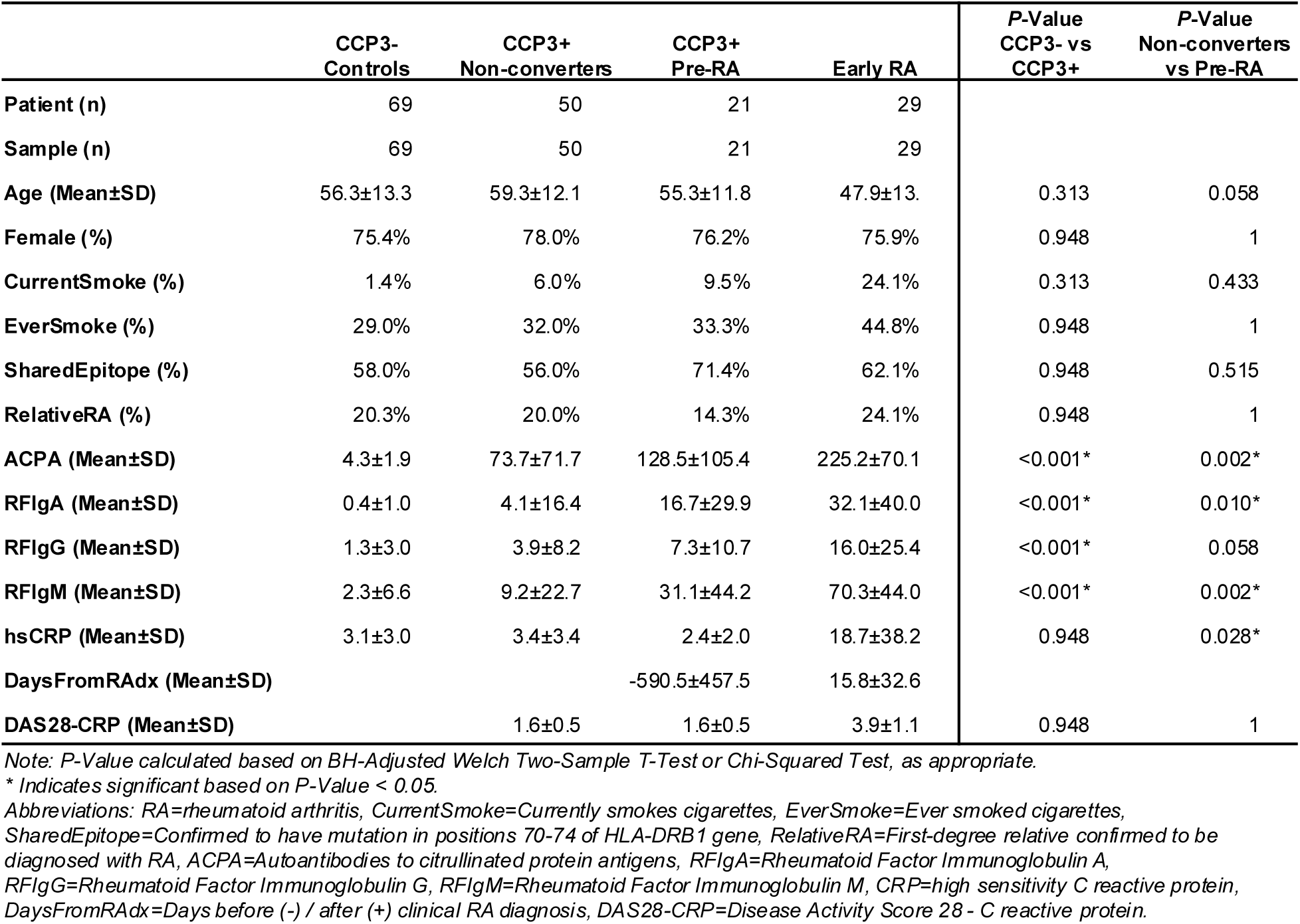
Baseline Participant Information.

|  | CCP3-<br>Controls | CCP3+<br>Non-converters | CCP3+<br>Pre-RA | Early RA | P-Value<br>CCP3- vs<br>CCP3+ | P-Value<br>Non-converters<br>vs Pre-RA |
| --- | --- | --- | --- | --- | --- | --- |
| Patient (n) | 69 | 50 | 21 | 29 |  |  |
| Sample (n) | 69 | 50 | 21 | 29 |  |  |
| Age (Mean±SD) | 56.3±13.3 | 59.3±12.1 | 55.3±11.8 | 47.9±13. | 0.313 | 0.058 |
| Female (%) | 75.4% | 78.0% | 76.2% | 75.9% | 0.948 | 1 |
| CurrentSmoke (%) | 1.4% | 6.0% | 9.5% | 24.1% | 0.313 | 0.433 |
| EverSmoke (%) | 29.0% | 32.0% | 33.3% | 44.8% | 0.948 | 1 |
| SharedEpitope (%) | 58.0% | 56.0% | 71.4% | 62.1% | 0.948 | 0.515 |
| RelativeRA (%) | 20.3% | 20.0% | 14.3% | 24.1% | 0.948 | 1 |
| ACPA (Mean±SD) | 4.3±1.9 | 73.7±71.7 | 128.5±105.4 | 225.2±70.1 | <0.001* | 0.002* |
| RFlgA (Mean±SD) | 0.4±1.0 | 4.1±16.4 | 16.7±29.9 | 32.1±40.0 | <0.001* | 0.010* |
| RFlgG (Mean±SD) | 1.3±3.0 | 3.9±8.2 | 7.3±10.7 | 16.0±25.4 | <0.001* | 0.058 |
| RFlgM (Mean±SD) | 2.3±6.6 | 9.2±22.7 | 31.1±44.2 | 70.3±44.0 | <0.001* | 0.002* |
| hsCRP (Mean±SD) | 3.1±3.0 | 3.4±3.4 | 2.4±2.0 | 18.7±38.2 | 0.948 | 0.028* |
| DaysFromRADx (Mean±SD) |  |  | -590.5±457.5 | 15.8±32.6 |  |  |
| DAS28-CRP (Mean±SD) |  | 1.6±0.5 | 1.6±0.5 | 3.9±1.1 | 0.948 | 1 |

Note: P-Value calculated based on BH-Adjusted Welch Two-Sample T-Test or Chi-Squared Test, as appropriate.
\* Indicates significant based on P-Value < 0.05.
Abbreviations: RA=rheumatoid arthritis, CurrentSmoke=Currently smokes cigarettes, EverSmoke=Ever smoked cigarettes, SharedEpitope=Confirmed to have mutation in positions 70-74 of HLA-DRB1 gene, RelativeRA=First-degree relative confirmed to be diagnosed with RA, ACPA=Autoantibodies to citrullinated protein antigens, RFlgA=Rheumatoid Factor Immunoglobulin A, RFlgG=Rheumatoid Factor Immunoglobulin G, RFlgM=Rheumatoid Factor Immunoglobulin M, CRP=high sensitivity C reactive protein, DaysFromRADx=Days before (-) / after (+) clinical RA diagnosis, DAS28-CRP=Disease Activity Score 28 - C reactive protein.

DNA methylation assays were performed in 3 batches, and these data were then combined. A batch effect was observed (row 3, column 2 of Supplementary Fig. 2) and was addressed using the Harman method of batch correction^8^. After correction, dimension reduction based on an unbiased analysis of all filtered autosomal loci readily distinguished B cells, CD4+ memory T cells and CD4+ naïve T cells, serving as an internal control on data reliability (Supplementary Fig. 3).

### Cross-Sectional Analysis Reveals Pre-Conversion Epigenetic Signature in “At-Risk” Participants at Baseline

We first expanded our previous cross-sectional analysis by performing pairwise comparisons among our participant samples at baseline (see Methods). With the additional knowledge indicating which anti-CCP3+ “At-Risk” participants would later convert to clinical RA within the study timeline, we could distinguish between anti-CCP3-“Controls”, anti-CCP3+ “Non-converters”, anti-CCP3+ “Pre-RA” and “Early RA” groups (Table 1). Baseline anti-CCP3+ Pre-RA samples were collected a mean of 591±458 (mean<u>+SD)</u> days before the time of first identification of clinical RA.

A pairwise comparison based on clinical status (Table 2) showed the greatest methylation differences between the anti-CCP3+ Pre-RA and RA groups in all cell lineages (360, 276, and 292 DMLs in CD19+ B cell, memory CD4+ T Cell, and naive CD4+ T cell samples, respectively). In addition, both of these groups exhibited large differences compared to all other clinical classifications (see Methods: DML, DMG & Pathway Analysis). In contrast, anti-CCP3-Controls and anti-CCP3+ Non-converters were similar to each other (only 3, 3, and 4 DMLs in B cell, CD4+ memory T Cell, and CD4+ naive T cell samples, respectively). Dimension reduction plots were generated using PCA (Fig. 1) based on the union of all DMLs found in comparisons of clinical status within each cell lineage. The first dimension highlights the differences between Pre-RA and Early RA while the second separates Pre-RA and Early RA samples from anti-CCP3+ Non-converters and anti-CCP3-Controls (see Methods: Chromatin State Analysis). We observed minimal differences between anti-CCP3-Control samples and anti-CCP3+ Non-converter samples while all other clinical classifications could be clearly distinguished from each other, consistent with the quantity of DMLs (Table 2).

**Figure 1.**
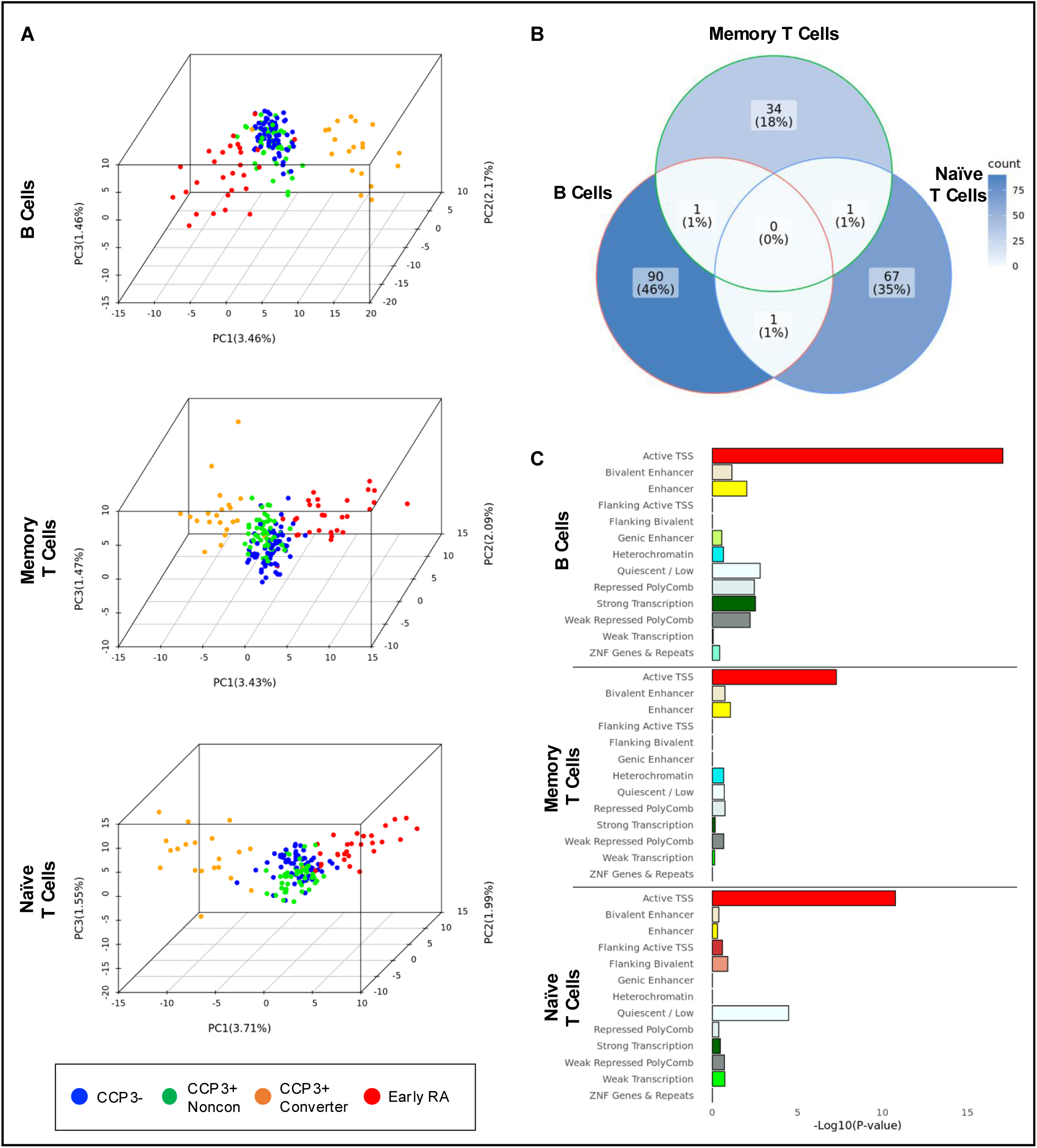
Cross-Sectional Analysis of Individuals at Baseline. (A) PCA of all baseline samples based on the union of DMLs derived from a pairwise comparison of clinical status including CCP3-Controls (n=69), CCP3+ Nonconverters (n=50), CCP3+ Pre-RA (n=21), and Early RA (n=29). (B) Venn diagram of the DMGs identified in CCP3+ Converters and Nonconverters. (C) Chromatin state enrichment within DMLs found between Pre-RA and Nonconverter participants at baseline. *Abbreviations: PCA=principal components analysis, DML=differentially methylated locus, DMG=differentially methylated gene, FDR=false discovery rate*.

**Table 2.**
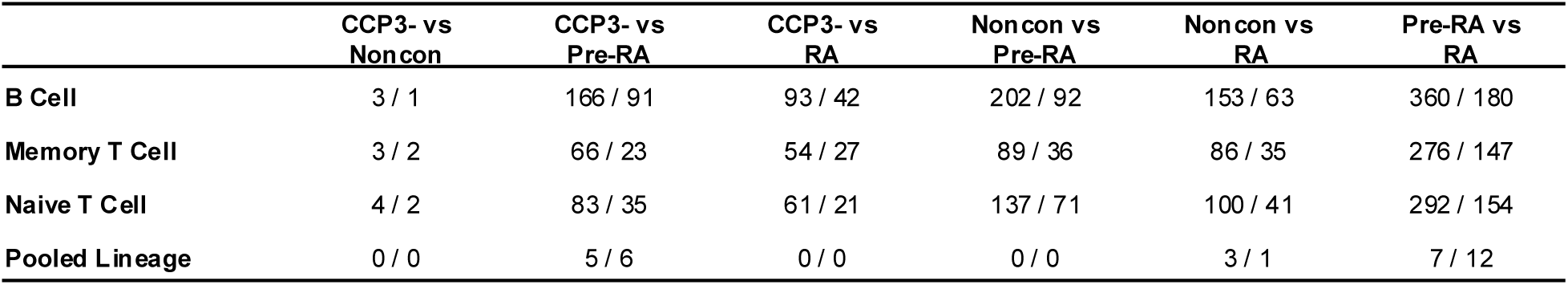
The number of DMLs / DMGs identified in cross sectional analysis among CCP3-controls, CCP3+ Nonconverter, CCP3+ Pre-RA, and Early RA in B cell, memory T cell, naïve T cell, and pooled lineage (in DMLs/DMGs), based on P-Value < 0.05, Difference of M > 2.0.

The DML overlap of our data with a previous study^9^ was limited, most likely because of our stringent filters. Therefore, we generated an additional set of DMLs using less stringent criteria in order to compare results for Early RA (Supplementary Table 1). The DMLs that we identified in T lymphocytes based on Early RA and Control data significantly overlapped with DNA methylome changes in Early RA (p = 3.55 × 10^-92^, based on union of CD4+ memory and naïve T cell DMLs by hypergeometric test)^9^.

We were particularly interested in identifying the baseline differences between anti-CCP3+ participants who would not convert to RA and those who would later progress to clinical RA. Note that the greatest differences in methylation between these groups occurred in B cells, followed by CD4+ naive T cells and CD4+ memory T cells (202, 137, and 89 DMLs, respectively). All lineages were enriched for Active Transcription Start Site (Active TSS). B cell DMLs from this comparison were additionally significantly enriched for Enhancer, Strong Transcription, Repressed Polycomb, Weak Repressed Polycomb and Epigenetically Quiescent chromatin states. CD4+ naive T cells were also enriched for Epigenetically Quiescent chromatin (Fig. 1).

We mapped the loci from each comparison to genes and plotted a Venn diagram to visualize the differentially methylated genes (DMGs) identified in the comparison of anti-CCP3+ Non-converter and Pre-RA participants at baseline (Fig. 1). Interestingly, in these analyses there was almost no overlap between DMGs across cell lineages. We then examined whether the distribution of DMGs suggested stochastic differences between Pre-RA and Non-converter participants or whether they aligned with known pathways. Pathway enrichment analysis of DMGs associated with CD4+ memory T cells and CD4+ naive T cells failed to identify significant pathways (FDR < 0.10). However, we identified 17 significantly enriched pathways based on DMGs associated with B cell samples (Supplementary Table 2). The enriched pathways were related to NOTCH signaling and nucleotide excision repair (each of which have been associated with RA^10–13^, suggesting a non-random methylation trajectory that distinguishes between Pre-RA and anti-CCP3+ Non-converter participants prior to disease onset.

### Methylome Remodeling in “At-Risk” Individuals Before RA Conversion

We next evaluated whether there was evidence of differential methylome remodeling from baseline over time for anti-CCP3-controls, anti-CCP3+ Non-converter participants and Pre-RA prior to conversion. Participants with baseline and year 1 samples in anti-CCP3-Control, anti-CCP3+ Non-converter and Pre-RA groups were analyzed (Table 3). Pre-RA participants were limited to those who had not converted to clinical RA after 1 year (n=7, Supplementary Fig. 1). Samples from Pre-RA individuals in this comparison were collected 714 ± 191 and 374 ± 177 days before clinical RA diagnosis for the baseline visit and the year 1 visit, respectively. During this period, no significant changes were observed for anti-CCP3, hsCRP or IgM, IgG, and IgA RF for either group (*p* < 0.05, BH-adjusted Welch Two Sample t-test and Chi-Square test).

**Table 3.**
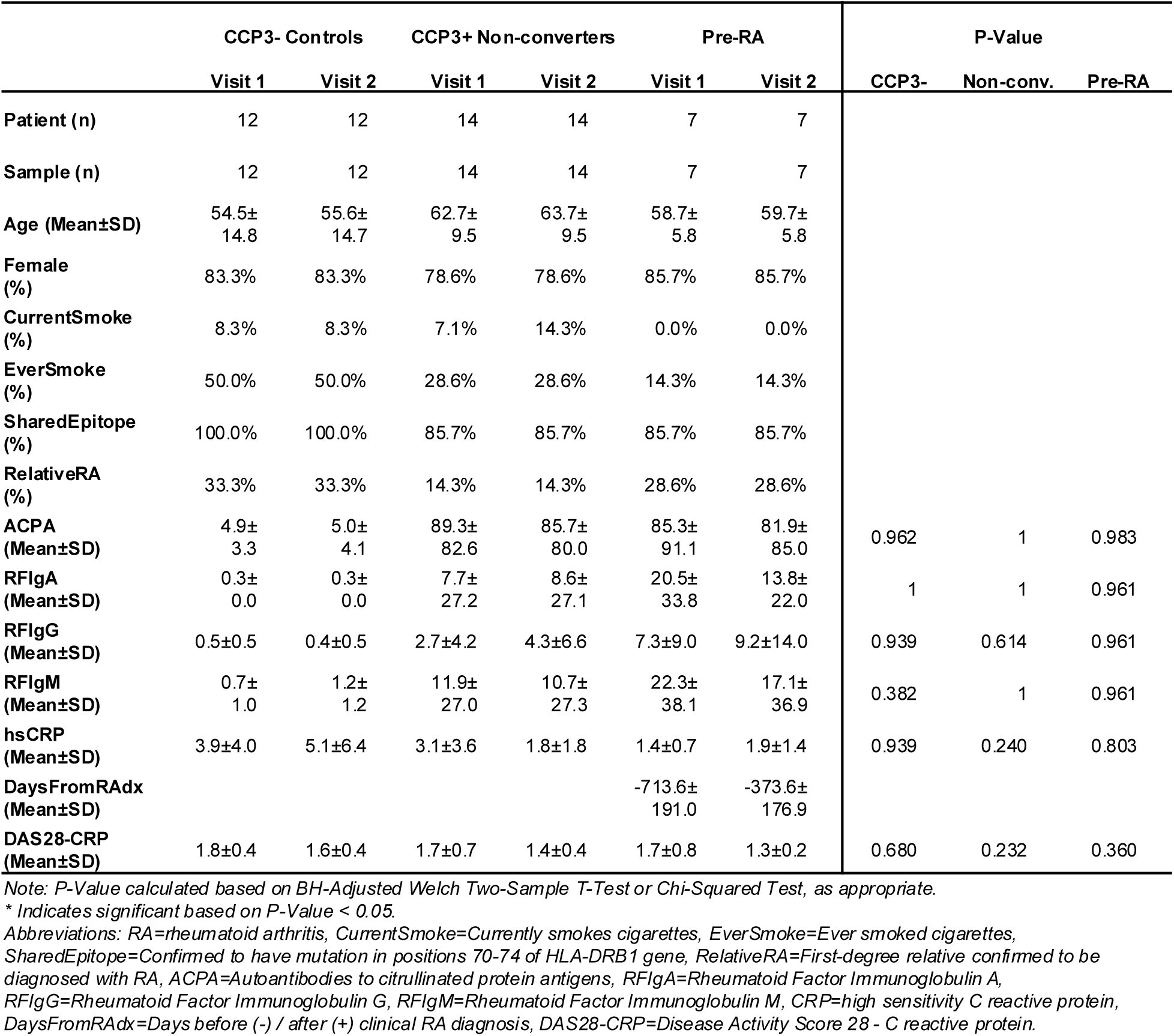
Paired Year 1 Longitudinal Comparison Participant Information.

|  | CCP3- Controls |  | CCP3+ Non-converters |  | Pre-RA |  | P-Value |  |  |
| --- | --- | --- | --- | --- | --- | --- | --- | --- | --- |
|  | Visit 1 | Visit 2 | Visit 1 | Visit 2 | Visit 1 | Visit 2 | CCP3- | Non-conv. | Pre-RA |
| Patient (n) | 12 | 12 | 14 | 14 | 7 | 7 |  |  |  |
| Sample (n) | 12 | 12 | 14 | 14 | 7 | 7 |  |  |  |
| Age (Mean±SD) | 54.5±<br>14.8 | 55.6±<br>14.7 | 62.7±<br>9.5 | 63.7±<br>9.5 | 58.7±<br>5.8 | 59.7±<br>5.8 |  |  |  |
| Female (%) | 83.3% | 83.3% | 78.6% | 78.6% | 85.7% | 85.7% |  |  |  |
| CurrentSmoke (%) | 8.3% | 8.3% | 7.1% | 14.3% | 0.0% | 0.0% |  |  |  |
| EverSmoke (%) | 50.0% | 50.0% | 28.6% | 28.6% | 14.3% | 14.3% |  |  |  |
| SharedEpitope (%) | 100.0% | 100.0% | 85.7% | 85.7% | 85.7% | 85.7% |  |  |  |
| RelativeRA (%) | 33.3% | 33.3% | 14.3% | 14.3% | 28.6% | 28.6% |  |  |  |
| ACPA (Mean±SD) | 4.9±<br>3.3 | 5.0±<br>4.1 | 89.3±<br>82.6 | 85.7±<br>80.0 | 85.3±<br>91.1 | 81.9±<br>85.0 | 0.962 | 1 | 0.983 |
| RFlgA (Mean±SD) | 0.3±<br>0.0 | 0.3±<br>0.0 | 7.7±<br>27.2 | 8.6±<br>27.1 | 20.5±<br>33.8 | 13.8±<br>22.0 | 1 | 1 | 0.961 |
| RFlgG (Mean±SD) | 0.5±0.5 | 0.4±0.5 | 2.7±4.2 | 4.3±6.6 | 7.3±9.0 | 9.2±14.0 | 0.939 | 0.614 | 0.961 |
| RFlgM (Mean±SD) | 0.7±<br>1.0 | 1.2±<br>1.2 | 11.9±<br>27.0 | 10.7±<br>27.3 | 22.3±<br>38.1 | 17.1±<br>36.9 | 0.382 | 1 | 0.961 |
| hsCRP (Mean±SD) | 3.9±4.0 | 5.1±6.4 | 3.1±3.6 | 1.8±1.8 | 1.4±0.7 | 1.9±1.4 | 0.939 | 0.240 | 0.803 |
| DaysFromRAdx (Mean±SD) |  |  |  |  | -713.6±<br>191.0 | -373.6±<br>176.9 |  |  |  |
| DAS28-CRP (Mean±SD) | 1.8±0.4 | 1.6±0.4 | 1.7±0.7 | 1.4±0.4 | 1.7±0.8 | 1.3±0.2 | 0.680 | 0.232 | 0.360 |

Note: P-Value calculated based on BH-Adjusted Welch Two-Sample T-Test or Chi-Squared Test, as appropriate.
\* Indicates significant based on P-Value < 0.05.
Abbreviations: RA=rheumatoid arthritis, CurrentSmoke=Currently smokes cigarettes, EverSmoke=Ever smoked cigarettes, SharedEpitope=Confirmed to have mutation in positions 70-74 of HLA-DRB1 gene, RelativeRA=First-degree relative confirmed to be diagnosed with RA, ACPA=Autoantibodies to citrullinated protein antigens, RFlgA=Rheumatoid Factor Immunoglobulin A, RFlgG=Rheumatoid Factor Immunoglobulin G, RFlgM=Rheumatoid Factor Immunoglobulin M, CRP=high sensitivity C reactive protein, DaysFromRAdx=Days before (-) / after (+) clinical RA diagnosis, DAS28-CRP=Disease Activity Score 28 - C reactive protein.

A paired analysis within clinical classifications was performed by comparing each participant’s baseline and year 1 samples in each cell lineage. There were minimal changes in methylation after one year for anti-CCP3-Control and anti-CCP3+ Non-converter individuals. Despite the limited number of participants, Pre-RA participants demonstrated significant methylome remodeling over the course of one year (162, 199, and 186 DMLs, for B cell, CD4+ memory T cell and CD4+ naive T cell samples, respectively) (Fig. 2 and Table 4).

**Figure 2.**
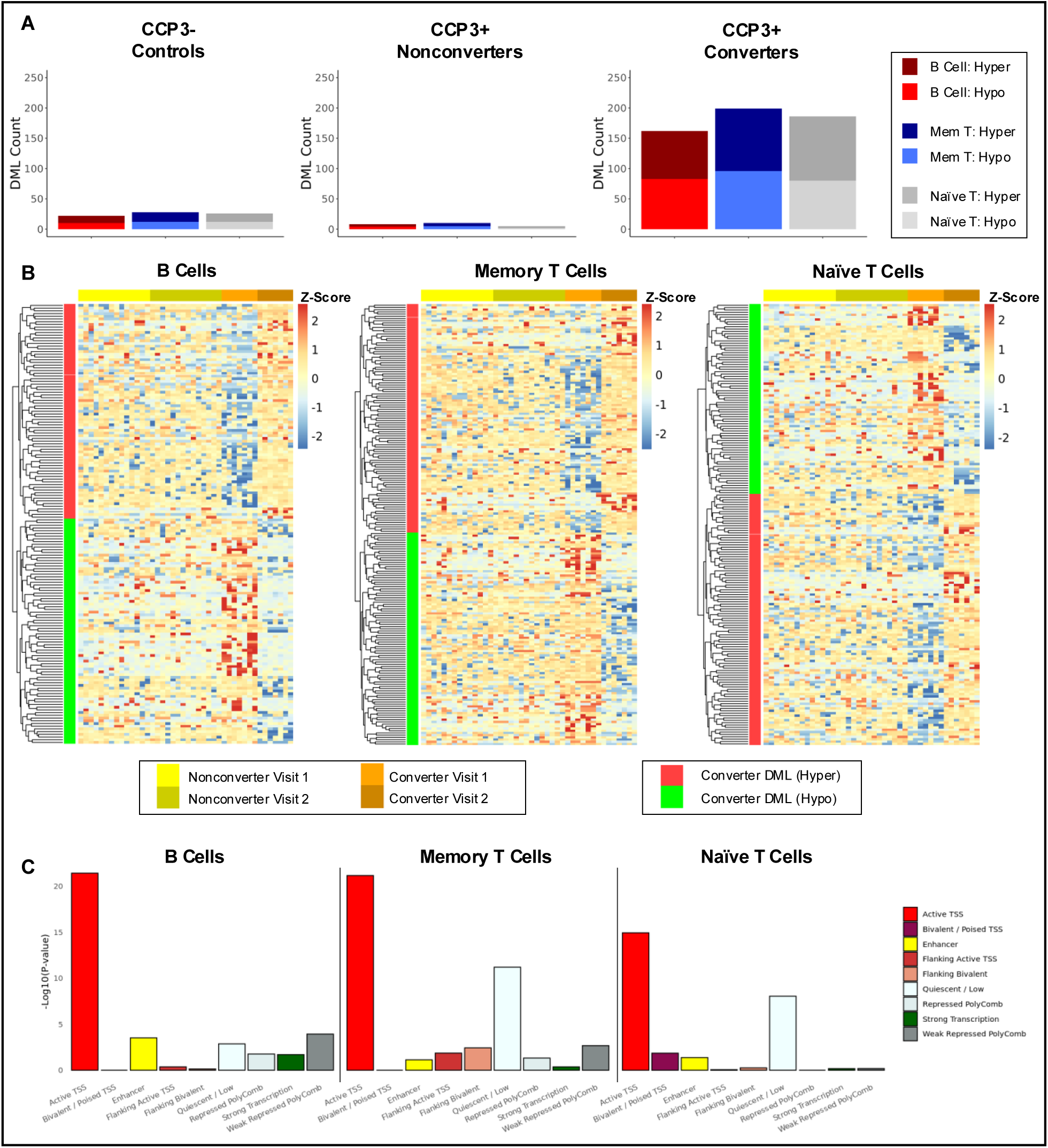
Methylome Remodeling in Early-Stage Pre-RA Individuals. (A) DML counts of CCP3-Controls (n=12), CCP3+ Nonconverters (n=14), and CCP3+ Pre-RA (n=7) between baseline and visit 2. (B) Heatmaps with rows showing DML found between Pre -RA participants baseline and year 1 visits and columns showing assays from CCP3+ Nonconverters and Pre-RA at baseline and visit 2. (C) Chromatin state enrichment within DMLs found between Pre-RA participants baseline and year 1 visits. *Abbreviations: DML=differentially methylated locus; DMG=differentially methylated genes*.

**Table 4.** The number of DMLs / DMGs identified in paired longitudinal analysis among CCP3 - controls, CCP3+ Nonconverter, and CCP3+ Converter, in B cell, memory T cell, and naïve T cell samples (in DMLs/DMGs), based on P-Value < 0.05, Difference of M > 3.5.

|  | CCP3-<br>Control | CCP3+<br>Nonconverter | CCP3+<br>Pre-RA |
| --- | --- | --- | --- |
| B Cell | 22 / 7 | 8 / 5 | 162 / 65 |
| Memory T Cell | 28 / 13 | 10 / 5 | 199 / 80 |
| Naïve T Cell | 26 / 12 | 5 / 1 | 186 / 72 |

Heatmaps of anti-CCP3+ Non-converter and Pre-RA samples based on identified Pre-RA DMLs through year 1 demonstrated clear separation of baseline samples from year 1 samples in Pre-RA patients within each cell lineage (Fig. 2). DMLs were mapped to genes and DMGs were used to perform pathway enrichment analysis. No pathways from any cell lineage met the enrichment significance threshold, but the identified DMLs from each cell lineage were significantly enriched for Active TSS while CD4+ memory T cell and CD4+ naïve T cell DMLs were also enriched for epigenetically quiescent regions (Fig. 2). Taken together, these results suggest that a trajectory of epigenetic modifications occur in at-risk pre-RA individuals that ultimately develop RA, and that these changes occur in regions highly relevant to transcriptional regulation.

### DML Clusters Exhibit Differing Patterns of Methylome Remodeling Pre-And-Post Diagnosis

To explore methylome remodeling both prior to and after the development of clinical RA, we next examined DNA methylation patterns in Pre-RA participants as they progressed to RA conversion and then to the early stages of RA. Pre-RA participants who had a sample taken before conversion (471 ± 159 days before diagnosis), at conversion, and after conversion (176 ± 91 days after diagnosis were selected for analysis (n=6). Within this cohort, no autoantibody or acute phase reactant levels significantly changed from baseline to the time of diagnosis of clinical RA or from diagnosis to a post-diagnosis visit 1 (*p* < 0.05, BH-adjusted Welch Two Sample t-test). DAS28-CRP significantly increased from baseline to diagnosis (p = 7.13 × 10^-5^, BH-adjusted Welch Two Sample t-test); however, we observed a decreasing trend from diagnosis to post-diagnosis visit 1 (Table 5). Additionally, two patients began treatment between initial diagnosis and post-diagnosis visit 1, one of which received methotrexate and one received low dose prednisone.

**Table 5.**
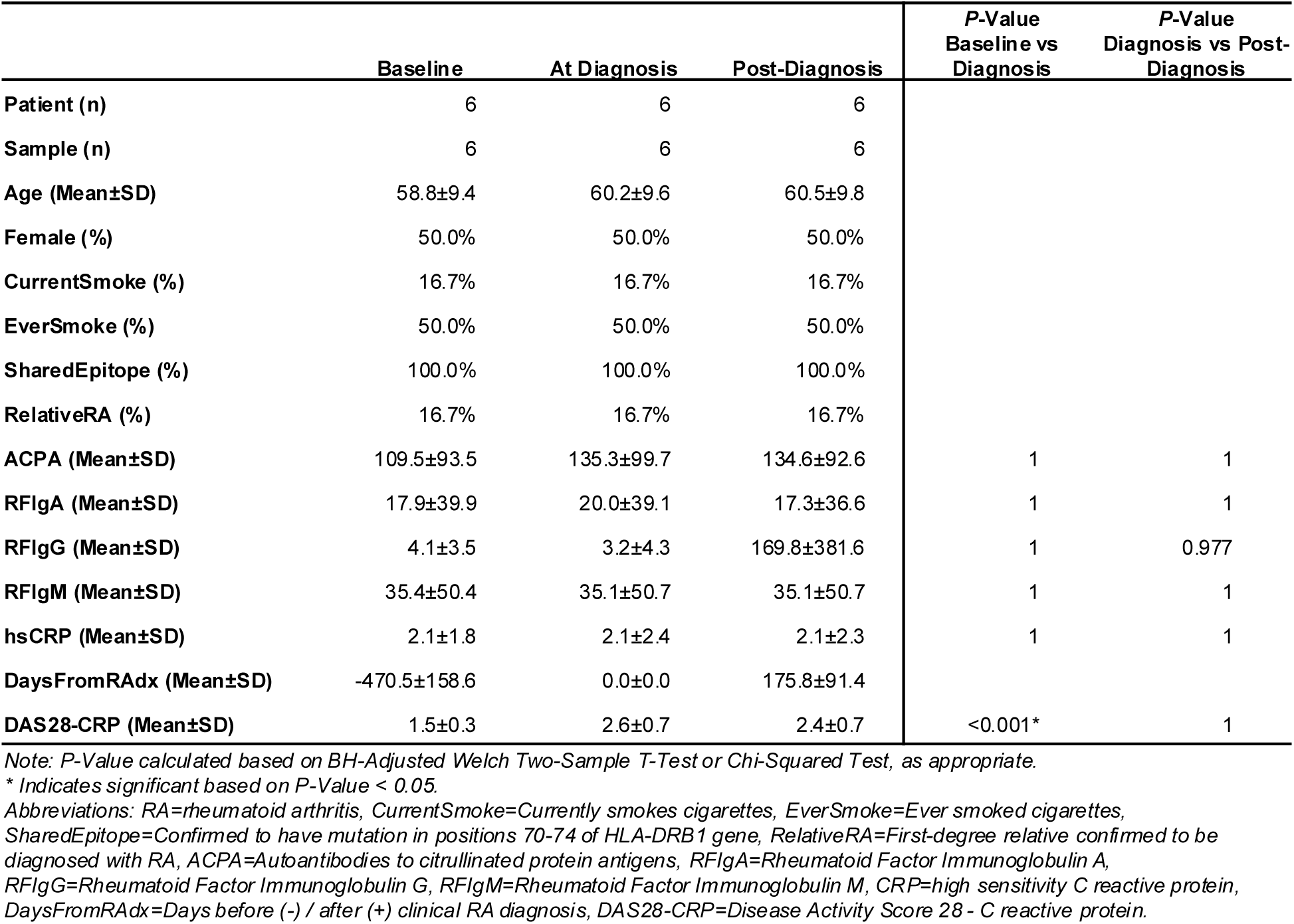
Paired Longitudinal Multi-Phase Comparison Participant Information.

A paired analysis of pre-diagnosis to diagnosis samples, diagnosis to post-diagnosis samples and pre-diagnosis to post-diagnosis samples was performed for each patient. DMLs were identified in each cell lineage. Similar numbers of DMLs were noted in each comparison with minimal overlap between cell lineages (Table 6).

**Table 6.** The number of DMLs / DMGs identified in paired longitudinal analysis among CCP3+ CCP3+ Pre-RA samples at multiple points in time, in B cell, memory T cell, and naïve T cell samples (in DMLs/DMGs), based on P-Value < 0.05, Difference of M > 3.5.

|  | Baseline to Dx | Dx to PostDx | Baseline to PostDx |
| --- | --- | --- | --- |
| B Cell | 290 / 124 | 241 / 104 | 326 / 148 |
| Memory T Cell | 291 / 137 | 233 / 101 | 261 / 113 |
| Naive T Cell | 332 / 154 | 246 / 114 | 341 / 159 |

We then determined the degree to which DML methylation changes were consistent between comparisons. Heatmaps of DMLs from each comparison for each cell lineage revealed multiple distinct patterns of DML remodeling (Fig. 3 and Supplementary Fig. 4). DMLs were then assigned to groups based on hierarchical clustering, with the optimal group number for each cell lineage selected using the elbow method (7, 8 and 8 groups for B cells, CD4+ memory T cells and CD4+ naïve T cells, respectively), and DMGs for each group were identified. Pathway enrichment analysis was performed on the union of all DMGs across clusters for each cell type (Fig. 3, Supplementary Tables 3-5). Two pathways were identified for CD4+ naive T cell samples, both related to caspase-signaling mediated apoptosis, but no significant pathways were found for B cells or CD4+ memory T cells (Supplementary Tables 3-5).

**Figure 3.**
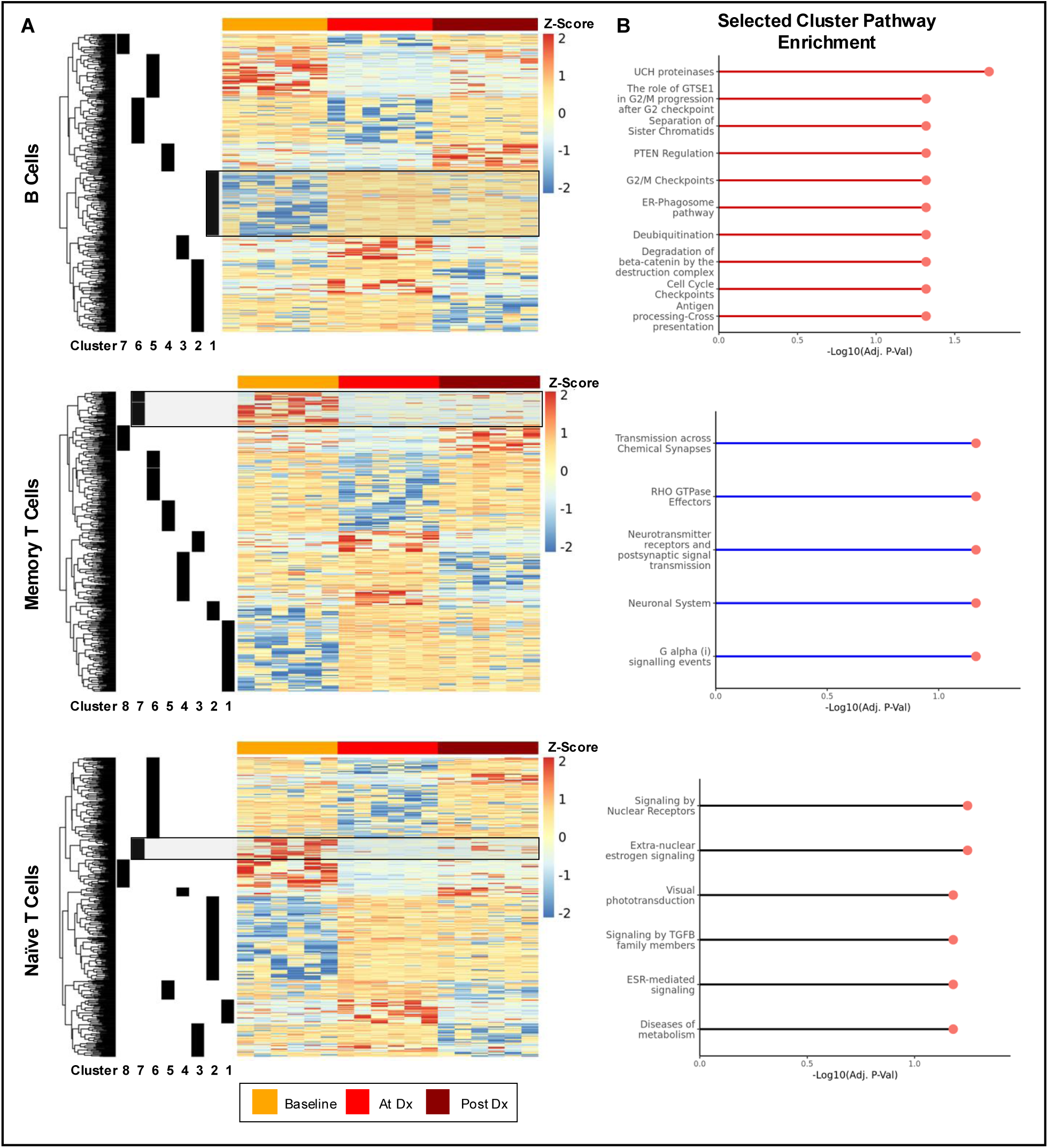
DML clusters exhibit multiple patterns of remodeling through RA progression. (A) Heatmaps with rows showing DMLs found in paired longitudinal analysis between CCP3+ Converters baseline, at-diagnosis, and post-diagnosis visits and columns showing assays from CCP3+ Converters baseline, at-diagnosis, and post-diagnosis visits (n=6). DML clusters are based on hierarchical clustering with cluster *k* selected using elbow method. Shading indicates DMLs associated with selected cluster for pathway enrichment. (B) Significantly enriched pathways found in comparison of individual clusters within B cells, memory T cells, and naïve T cells (FDR < 0.10). *Abbreviations: DML=differentially methylated locus; DMG=differentially methylated genes, FDR=false discovery rate*.

Because of the heterogeneity of DML methylation patterns in each cluster, we sought to understand the potential pathways enriched in individual DML clusters (Fig. 3). This analysis yielded more substantive pathway enrichment. Within B cells, cluster 1 DMGs were enriched for cell-cycle transition, protein degradation^14,15^, and adaptive immune system pathways. CD4+ memory T cell cluster 7 was enriched for synaptic transmission and signaling pathways. Within CD4+ naive T cells, cluster 7 was characterized by enrichment of nuclear signaling receptor pathways, including by TGFβ.

These clusters exhibit differing patterns of methylome remodeling during their trajectory before and after conversion to clinical RA and demonstrate the dynamic nature of these changes. For example, B cell cluster 1 DMLs were relatively hypomethylated at baseline, transitioned to a hypermethylated state at the time of diagnosis, and remained hypermethylated through early RA. C4+ naïve and memory T cell cluster 7 DMLs begin relatively hypermethylated before transitioning to a hypomethylated state at diagnosis and in early RA. Many DML clusters (e.g., B cell cluster 6, CD4+ memory T cell cluster 6, CD4+ naïve T cell cluster 6) were characterized by a different pattern where loci were hypomethylated when transitioning between pre-to-at-diagnosis and then later reverted to baseline hypermethylation when transitioning between diagnosis to post-diagnosis. Notably, clusters resulting in the most significant enriched pathways followed a monotonic trajectory.

### Epigenetic Markers Demonstrate Improved Prognostic Capability Compared to Auto-antibody Data

Having noted the epigenetic differences exhibited between Pre-RA and Non-converter samples at baseline and 1 year visits (Table 3, Figure 2), we next investigated whether specific epigenetic loci in the former are associated with RA conversion. To establish a performance baseline, we first built a classifier based only on autoantibody/acute phase reactant data to identify anti-CCP3+ participants at their initial visit that would go on to convert to clinical RA. These participants (n=71) were separated into training (70%) and test (30%) sets^16^ and autoantibody/acute phase reactant predictors (anti-CCP3, hsCRP, and IgM, IgG, and IgA rheumatoid factor) were ranked by importance using recursive feature elimination (RFE). After eliminating unimportant predictors (importance < 0, based on RFE algorithm), a random forest classification model was trained using 10-fold cross-validation, which was repeated 5 times. This process was iterated 100 times using different training and test splits at each predictor count, with models evaluated for performance based on training accuracy in each iteration (Fig. 4 and Supplementary Fig. 5). Using only continuous autoantibody and acute phase reactant data, the optimal predictor count was selected. This optimized prediction model achieved a training accuracy of 75.0% ± 5.1% and test accuracy of 71.6% ± 7.2% in predicting which baseline anti-CCP3+ patients would go on to convert to clinical RA. This model achieved Area Under the Curve (AUC) of 64.0% ± 10.8% and Positive Predictive Value (PPV) of 75.7% ± 5.6% on test data (all data provided as mean ± SD).

**Figure 4.**
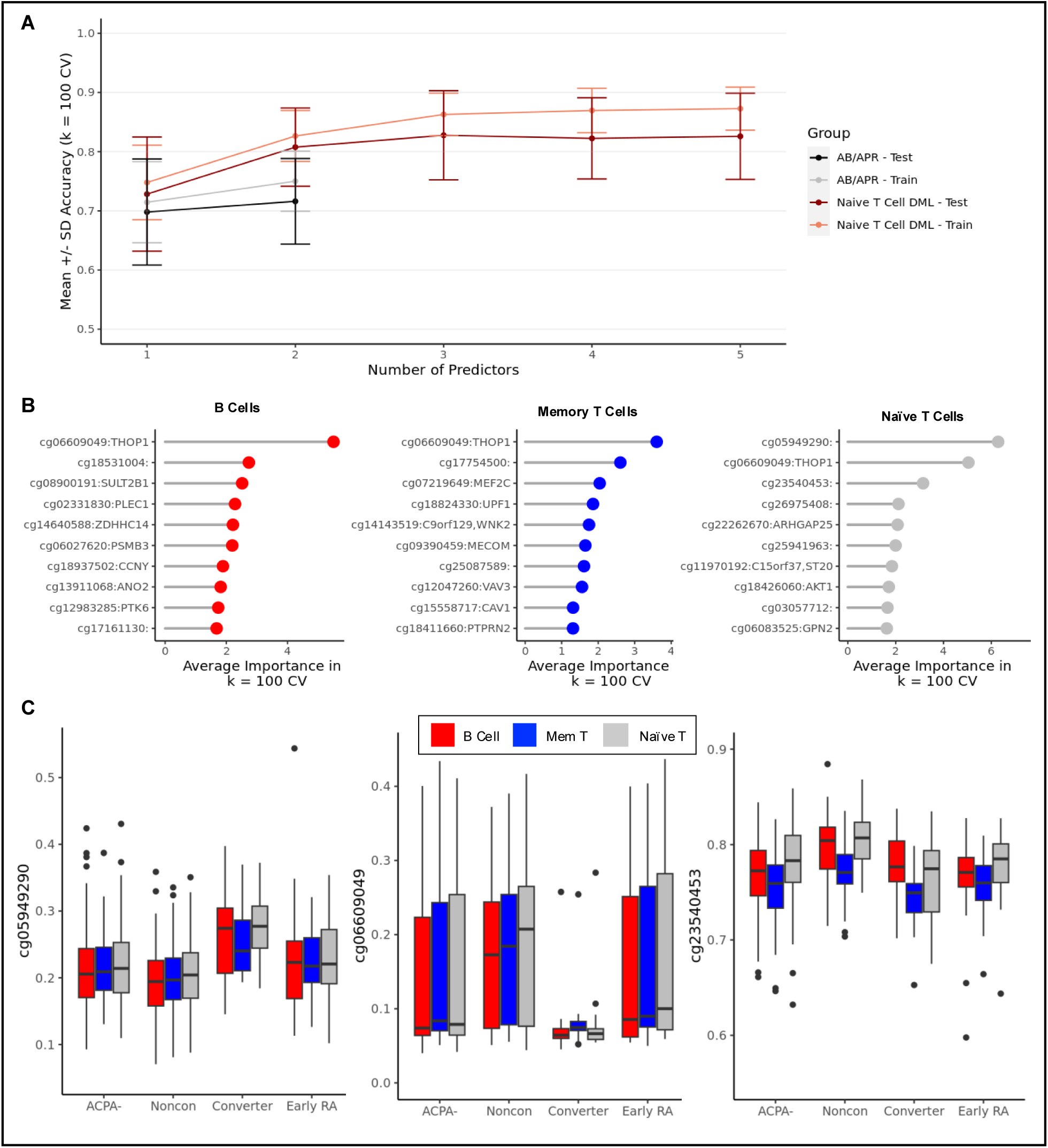
Predictive model identifies key risk loci for CCP3+ conversion. (A) Mean accuracy +/- standard deviation of predictive models from naïve T cell DMLs and autoantibodies. Accuracies were plotted up to the optimal training accuracy for each group. (B) Mean importance of DML predictors in B cells, memory T cells and naïve T cells. Importance metrics were calculated based on 100 iterations of data partitions and recursive feature elimination. (C) Methylation intensities (depicted as beta values) of the three most important DML predictors models derived from naïve T cells in baseline visit samples. *Abbreviations: DML=differentially methylated locus; DMG=differentially methylated genes*.

We then repeated this process using the methylation data from each cell lineage (Fig. 4 and Supplementary Fig. 5). After feature selection (see Methods), we trained random forest classifiers using increasing numbers of selected DMLs and selected the optimal predictor count for each cell lineage. Using B cell data, the prediction model achieved 84.0% ± 3.4% training accuracy and 73.7% ± 7.4% test accuracy using 7 predictors. The optimal classifier using memory CD4+ T cell data also utilized 4 predictors and resulted in 82.6% ± 3.8% training accuracy and 71.8% ± 8.3% test accuracy. The model utilizing naive CD4+ T cell data was the most accurate, achieving 87.3% ± 3.6% training accuracy and 82.6% ± 7.3% test accuracy while using 5 predictors (Table 7). This model also achieved an AUC of 87.2% ± 9.0% and PPV of 84.5% ± 6.3% on test data. Models utilizing methylation markers outperformed the optimal baseline autoantibody/acute phase reactant classifier in all cell lineages, and improvement observed in the predictive model derived from CD4+ naive T cells was highly significant (*p* = 2.20 × 10^-16^, Welch Two Sample t-test). Combining DML and autoantibody/acute phase reactant data resulted in slightly higher training accuracies for each lineage, but no increase in test accuracies. Additionally, combining methylation data from each cell lineage into a single integrated model similarly increased training accuracy (88.1% ± 3.2%), with no resulting increase in test accuracy over the CD4+ naïve T cell model (77.1% ± 7.6%). This could indicate that these datasets are not additive or that this sample set was underpowered to take advantage of the larger feature set.

**Table 7.** Comparison of Random Forest model performance metrics. Data shown represents values derived from model using parameters yielding highest training accuracy.

|  | Predictors | Training Accuracy | Test Accuracy | Test AUC | Test PPV |
| --- | --- | --- | --- | --- | --- |
| **B Cell** | 7 | 84.0% ± 3.4% | 73.7% ± 7.4% | 76.9% ± 9.9% | 77.8% ± 6.6% |
| **Memory T Cell** | 4 | 82.6% ± 3.8% | 71.8% ± 8.3% | 71.6% ± 10.8% | 77.5% ± 6.5% |
| **Naïve T Cell** | 5 | 87.3% ± 3.6% | 82.6% ± 7.3% | 87.2% ± 9.0% | 84.5% ± 6.3% |
| **Cell Lineage Combined** | 9 | 88.1% ± 3.2% | 77.1% ± 7.6% | 83.3% ± 7.7% | 84.0% ± 7.2% |
| **B Cell + AB/APR Data** | 9 | 85.2% ± 3.5% | 73.8% ± 7.6% | 77.3% ± 10.5% | 81.7% ± 6.8% |
| **Memory T Cell + AB/APR Data** | 7 | 83.8% ± 3.4% | 72.0% ± 7.5% | 71.1% ± 10.7% | 76.5% ± 6.2% |
| **Naïve T Cell + AB/APR Data** | 5 | 87.3% ± 3.4% | 80.7% ± 7.2% | 86.6% ± 8.5% | 83.3% ± 6.9% |
| **AB/APR Data Only** | 2 | 75.0% ± 5.1% | 71.6% ± 7.2% | 64.0% ± 10.8% | 75.7% ± 5.6% |

*Note: Data show mean ± standard deviation over $k=100$ runs.*
*Abbreviations: AB=Antibody, APR=Acute Phase Reactant*

We next evaluated the key DMLs that drive the predictive power of our models. One DML, cg06609049, was in the top 10 features of all three cell lineages based on prediction importance (Fig. 4). cg06609049 is located at position chr19:2785107, a region flanking the active transcription start site in the promoter region of *THOP1* within CD4+ naive T cells. This location is significantly hypermethylated in Non-converters compared to Controls (*p* = 6.66 × 10^-2^, *p* = 8.13 × 10^-2^, and *p* = 9.95 × 10^-2^ for B cells, CD4+ memory T cells and CD4+ naive T cells, respectively, based on Welch Two Sample t-test) and in Pre-RA (*p* = 1.85 × 10^-5^, *p* = 6.43 × 10^-5^, and *p* = 4.78 × 10^-5^ for B cells, CD4+ memory T cells and CD4+ naive T cells, respectively, based on Welch Two Sample t-test), but not compared to Early RA (Fig. 4 and Supplementary Fig. 6).

Noting the increased predictive power of models using CD4+ naive T cell epigenetic markers, we also examined the most important probes contributing to the CD4+ naive T cell model, in addition to cg06609049 (Fig. 4). cg05949290 and cg23540453 were identified as the first and third most important predictors of conversion status, respectively. Neither are located in an annotated regulatory region. Further, the core 15-state model provided by the Roadmap Epigenomics Project indicates their locations are epigenetically quiescent in CD4+ naive T cells. cg06609049 is significantly hypermethylated in Non-converters compared to Pre-RA participants (*p* = 1.85 × 10^-5^, *p* = 6.43 × 10^-5^, and *p* = 4.78 × 10^-5^ for B cells, CD4+ memory T cells and CD4+ naive T cells, respectively, based on Welch Two Sample t-test). cg142957727 is most significantly hypermethylated in CD4+ naive T cells when comparing Non-converters and Pre-RA (*p* = 3.68 × 10^-6^), but it is also hypermethylated in B cells and CD4+ memory T cells to a lesser extent (*p* = 1.03 × 10^-2^ and *p* = 6.30 × 10^-4^, respectively).

## Discussion

To understand the trajectory of epigenetic marks in the journey from pre-RA to clinical synovitis, we performed a unique longitudinal analysis of anti-CCP3-Controls, anti-CCP3+ At-Risk individuals, and Early RA patients. Epigenetic remodeling occurred in individual participants over time as measured by DNA methylation in three PBMC-derived cell lineages that had been separated experimentally rather than by algorithmic deconvolution. DMLs were identified that distinguish the clinical cohorts and construction of machine learning models demonstrated that specific loci could predict which At-Risk individuals would later convert to clinical RA. This longitudinal analysis represents the first time that an epigenetic signature distinguishing anti-CCP3+ Pre-RA individuals from anti-CCP3+ Non-converters has been defined. Further, it marks the first observation of a dynamic “rheumatoid arthritis methylome trajectory” in Pre-RA. These findings help characterize important pathogenic events in the continuum of RA pathogenesis and hold potential for prognostic tests and identification of novel strategies to prevent RA.

Previous studies have characterized many aspects of the epigenetic contribution to RA pathogenesis, with many focusing on the methylation patterns of Early RA. For example, fibroblast-like synoviocytes (FLS) have distinctive methylation patterns in Early RA patients compared with FLS isolated from individuals with longstanding disease^17^. Additional research into FLS has identified genomic loci with methylation levels that correlated with disease severity in RA patients^18^. Within lymphocyte subpopulations derived from whole blood, others found that treatment naïve early RA patients display aberrant DNA methylation at specific loci^9^. A study of monocytes, naïve and memory CD4+ T cells found that early RA patients displayed altered methylation patterns within an IL6/JAK1/STAT3 node related to TNF-signaling and Th17 cell differentiation^19^.

Other studies identified epigenetic factors in early undifferentiated inflammatory arthritis (UA) prior to developing classifiable RA. For instance, one report identified differential methylation patterns in FLS derived from patients with early UA^20^. Similarly, an analysis of PBMCs from UA patients found a methylation signature associated with progression to RA^7^. Our previous cross-sectional analysis of the TIP-RA cohort also showed that peripheral blood lymphocytes of anti-CCP3+ “At-Risk” individuals were epigenetically distinct from Early RA patients as well as anti-CCP3-Controls^6^. These analyses provided initial insights into the immune dysregulation of anti-CCP3+ patients, with some potential relevance as individuals fully progress from asymptomatic autoimmunity to clinical RA. These studies focused on mixed populations of PBMCs, which could limit their power.

Although other cross-sectional data on PBMC DNA methylation in at-risk populations and RA are available in addition to our own TIP-RA studies^19,21–23^, the present analysis is unique in its ability to follow methylation remodeling through the clinical transitions in single individuals over time using purified T and B cell subsets. This study design allowed us to use highly stringent criteria to define DMLs and mitigate confounding influences like sex, cell mixtures, and the influence of SNPs on methylation. Using these filters, we found that our DMLs had limited overlap with previous datasets. However, significant overlap emerged when we decreased stringency, validating our results. Taken together, our data provides the first comprehensive map of the methylation trajectory through the natural history of RA, from asymptomatic autoimmunity through to clinical RA.

One of the most striking findings was the stability of methylomes on anti-CCP3-controls and anti-CCP3+ non-converters over time compared with remodeling in anti-CCP3+ converters both before and after the development of clinical RA. We are uncertain about the mechanisms behind these dynamic changes, some of which lead to increased and some of which lead to decreased methylation at individual loci. The modified loci are highly enriched in active promoter elements as well as epigenetically quiescent regions, indicating they could be the result of multiple processes. It is also unknown if these changes are directed or stochastic, whether they occur throughout the blood, mucosal surfaces and central lymphoid organs, or whether similar changes are occurring within or around the joint tissues.

Numerous environmental factors can contribute to methylome remodeling, and we currently cannot determine which are responsible for the changes or serve as triggers for the epigenetic inflection towards RA. Demographic factors like age^24^ and sex^25^, as well as other exposures like smoking status^26,27^, are known to influence epigenetic profiles. Our analysis has accounted for these factors as covariates in our baseline cross-sectional identification of DMLs. Not only that, there are minimal differences in these parameters across the study groups, suggesting that the answer might lie elsewhere.

An alternative explanation is that the earliest break in tolerance to self-antigens provides the catalyst for the initiation of an inflammatory milieu^4^ which drives methylome remodeling in circulating lymphocytes. For example, DNA methyltransferases (DNMTs) can deposit *de novo* 5mC modifications in response to this type of trigger^28^. The interaction of long non-coding RNAs (lncRNAs) and DNA methylation is one potential mechanism that can guide this process. This can occur both through methylation that regulates lncRNA expression^29^ or through lncRNAs recruiting DNMTs to certain loci^30^, which can regulate cytokine production^31^ within RA^32,33^. While the mechanism is not well characterized, *in vitro* exposure of healthy monocytes to cytokines can partially recapitulate the methylome of RA^34^. While this view of pathogenesis could explain the dynamic epigenetic trajectory we observe in Pre-RA, it still leaves unanswered the question of why some ACPA+ individuals remain epigenetically “static”, while others divert towards clinical RA.

We identified pathways enriched as autoimmunity progresses from pre-clinical to classifiable RA, but the number of pathways was unexpectedly low compared to the number of DMLs. Furthermore, no pathways were enriched in our analysis of Pre-RA individuals before conversion. This difference might be attributed to several factors, including the limitations of the technology used to identify DMLs. For example, the EPIC methylation chip, although focused on regulatory regions, covers only 27% of proximal and 7% of distal DNAse-sensitive sites identified by the ENCODE project^35^, which might restrict the ability to associate DMLs with genes and pathways. Furthermore, to mitigate sex-related differences, our study excluded the allosome, which is notably rich in immune-related genes^36,37^. It is also possible that several distinct processes drive the immune evolution, resulting in a more heterogenous pattern that obscures individual pathways in subsets. For example, changes in lymphocyte subset distributions have been observed in RA patients before and after treatment^38^, and differences are also present RA patients compared to controls^39^. Our study minimizes these influences by analyzing three experimentally-separated lineages. Still, it is possible that subtle composition differences within these separated lineages could have influenced our results. Further research utilizing single-cell, whole genome methylation sequencing is prohibitively expensive currently but could resolve this potential concern along with an expanded genomic focus. Incorporation of additional exposure and natural history data will be necessary to parse the degree to which these factors influenced our results. Despite these limitations, our ability to separate cells into at 3 separate lineages provides the greatest resolution for DNA methylation in at-risk and RA populations to date.

Notably, our analysis indicates that methylation changes driving NOTCH signaling in B cells were associated with clinical RA conversion in an anti-CCP3+ “At-Risk” population. Our findings point specifically to the role of NOTCH-1 signaling, but we note the significant overlap in genes, ligands, and function within the NOTCH family^40^, indicating the utility of broadly examining the role of NOTCH signaling in RA. NOTCH-3 signaling drives the differentiation of perivascular and sublining CD90(*THY1*)+ fibroblasts in synovial tissue organoids and contribute to inflammation in the rheumatoid joint^12^. *Notch3* targeting strategies, including knockout and antibody blockade, can ameliorate inflammation and bone damage in inflammatory arthritis^12^. NOTCH-1 signaling has also been implicated in RA pathogenesis, specifically by encouraging proliferation of TNF-activated FLS^10^. Further, *Notch1* has been identified as a possible drug target for RA. *In vivo* analysis demonstrates that *Notch1*-targeted siRNA nanoparticles can reduce progression of inflammation, bone erosion, and cartilage damage in a collagen-induced murine model^11^. Our findings build on this body of research, further highlighting the role of NOTCH signaling in RA progression.

With regards to our predictive biomarkers, cg06609049, a probe targeting the promoter region of *THOP1*, is strongly predictive of conversion in all three cell lineages. *THOP1* has a multi-faceted role in the MHC-I presentation pathway^41^ by positively^42^ or negatively^43^ regulating MHC-I antigen processing and expression^44^. Interestingly, *THOP1* expression is lower in RA compared with healthy controls in whole blood samples^45^. *THOP1* might contribute to RA pathogenesis and has been suggested as a potential target for autoimmune disease treatment^41^. Other key predictors driving model performance were found in epigenetically quiescent regions, suggesting that these loci could act through non-classical regulatory mechanisms or be the result, not the cause, of directed methylome remodeling associated with clinical conversion.

This investigation has not found a causative “epigenetic smoking gun” that determines which ACPA+ individuals will develop clinical RA, although the associations with NOTCH are intriguing. However, we do find evidence that Pre-RA individuals are epigenetically distinct from non-converters, which more closely resemble ACPA-controls. From this altered baseline, the Pre-RA methylome then continues to remodel on a dynamic trajectory towards clinical RA. Further research will be necessary to uncover the fundamental mechanisms that govern the early diversion of Pre-RA individuals from their phenotypically similar peers. However, these results do provide compelling evidence for the use of epigenetic biomarkers to predict the onset of clinical RA within an ACPA+ population, potentially increasing the precision and robustness with which clinical trials would be able to test novel strategies for prevention.

## Methods

### Sex as a Biological Variable

Our study examined both sexes and considered sex as a covariate. The study was not powered to investigate differences between sexes.

### Cohort and Biosamples

The Targeting Immune Responses for Prevention of Rheumatoid Arthritis (TIP-RA) cohort was designed to prospectively study individuals at high risk for developing RA due to the presence at baseline of serum anti-CCP3 positivity (IgG, Werfen, San Diego, CA) in the absence of a history of IA, or IA at a baseline physical examination of 66/68 joints. ACPA positive (anti-CCP3+) and ACPA negative (anti-CCP3)-subjects were recruited through screening of health-fair participants, first-degree relatives of patients with RA, and individuals referred for evaluation to rheumatology clinics. Individuals with RA were identified through rheumatology clinics and ACPA status was determined using the anti-CCP3 assay (IgG, Werfen, San Diego, CA). A detailed description of the study protocols for subject identification, clinical phenotyping, and clinical autoantibody testing has previously been described (see reference 6).

### Peripheral Blood Processing and DNA Methylation Measurement

Peripheral blood mononuclear cells were obtained from participants annually, processed and stored frozen in DMSO-based medium. Viability and yield were confirmed by dye-exclusion assay. Cell subsets were isolated by sequential magnetic separation using a Miltenyi Biotec AutoMacs Pro. CD19+, CD4+/CD45RO+ memory, CD4+/CD45RO-naive phenotypes were used for methylation analysis. Genomic DNA was isolated from 177 patients with anti-CCP3-, anti-CCP3+ and early RA with multiple visits. RNA-free PBMC subset genomic DNA was prepared by affinity column with DNeasy Blood & Tissue Kit (Qiagen) followed by concentration with Ultra-0.5 Centrifugal Filters (Amicon) and concentration normalized (20ng/µl). The Illumina Infinium MethylationEPIC Kit chip was used to measure DNA methylation levels at ∼850,000 CpG loci per sample.

### Beadchip Data Processing

Data generated using the Infinium MethylationEPIC Kit was processed using the minfi^46^ package v1.40.0 using a series of quality control, processing and normalization procedures. All processing and downstream analysis was performed using R version 4.2.3. Samples with a mismatch between described sex and reported sex using the minfi command getSex() were removed from analysis^47^. Samples were then separated into female and male groups and probe detection *p*-value analysis was performed^47^. Samples were re-combined and methylation signal intensity, beta outlier, bisulfite conversion and bead count quality control steps were performed.

Previous studies^48,49^ have described the potential bias that can arise from methylation analysis at CpG loci overlapping single nucleotide polymorphisms (SNPs), as well as probes that demonstrate cross-reactivity across multiple loci. To counter these potential sources of bias, we performed further quality control procedures. Previously identified probes overlapping common SNPs were removed from the analysis based on annotation in the IlluminaHumanMethylationEPICanno.ilm10b4.hg19^50^ R package using the minfi dropLociWithSnps() function. Further, we performed a targeted gap-hunting analysis using the MethylToSNP^51^ package v0.99.0 to eliminate probes with evidence of potential SNP overlap specifically within our data. DMLs identified as key predictors in the conversion prediction model were inspected to ensure methylation distributions were not consistent with SNP overlap (Supplementary Fig. 6). Finally, we removed potentially cross-reactive probes with > 47 bp homology, as previously described^35^.

Initial data processing and between-array normalization was performed using the preprocess Illumina method. After initial processing, principal components analysis (PCA) of our data indicated the presence of a batch effect. We used the Harman^8^ method of correction under default parameters to address this artifact (Supplementary Fig. 2 and 3). Harman minimizes batch noise by correcting DNA methylation measurements for batch effect under a probabilistic constraint for overcorrection. After correction, PCA was used to evaluate the effectiveness of batch effect removal.

Signal intensity values for each sample were extracted and calculated as both β and M-values. M-values were used in downstream statistical analysis and visualization due to β-value heteroscedasticity in high and low methylation ranges^52^, unless otherwise stated. Individual probe values were reported and visualized as β-values.

### DML, DMG & Pathway Analysis

Differentially Methylated Loci (DML) were identified using differences of mean M-value and empirical bayes-moderated t-test *p*-value criteria. Due to the limited number of DMLs identified, raw *p*-values were not adjusted for multiple testing corrections. All tests were performed using the limma^53^ R package version 3.50.3.

In our previous cross-sectional analysis^6^, the baseline sample count provided sufficient power to comprehensively investigate the entire genome while analyzing only female samples. For this analysis, we sought to identify the most important loci that could potentially act as predictive biomarkers in the broadest possible population. We thus applied highly stringent selection parameters utilizing sample from male and female participants while adjusting for important covariates (sex, age, and current smoking status). This methodology restricted the scope of our investigation to the autosome to avoid incongruent information between samples. *P*-value < 0.05 and mean |M| > 2.0 criteria were used for cross-sectional comparisons. All longitudinal analyses utilized *P*-value < 0.05 and mean |M| > 3.5 as DML selection criteria and were performed using paired samples from each patient.

To compare our results against data available in public databases, we also selected a DML set based on a cross-sectional analysis using less stringent criteria, consistent with previous investigations^9^. For this analysis DMLs were selected based on having P-Value < 0.05 and mean |M| > 0.5.

DMLs were then mapped to genes. Probe location on the GRCh37^54^ (hg19) assembly was determined using the IlluminaHumanMethylationEPICanno.ilm10b4.hg19 R package. GRCh37 gene body and promoter (TSS-2,500bp to TSS+500bp) coordinates were determined using the biomaRt^55^ R package version 2.50.3. Genes with a DML found in the gene body or promoter were determined to be differentially methylated genes (DMGs).

Enriched pathways were identified using the ReactomePA^56^ R package version 1.42.0 based on Reactome Database^57,58^ Schema version 82. Significantly enriched pathways were selected based on having an FDR < 0.1 (overrepresentation test, as calculated by ReactomePA), unless otherwise stated.

### Chromatin State Analysis

Chromatin state analysis was performed using cell lineage specific data (E032: Primary B cells, E040: Primary T helper memory cells, and E038: Primary T helper naive cells) from the core 15-state model provided by the Roadmap Epigenomics Project^59^. This core 15-state model was generated using ChromHMM^60^ on epigenetic data derived from 5 chromatin marks (H3K4me3, H3K4me1, H3K36me3, H3K27me3, H3K9me3) in 127 epigenomes. We performed enrichment analysis for chromatin states in each comparison within each cell lineage using a Fisher’s test based on all post-QC probes as a background. Significantly enriched (*p* < 0.05) chromatin states were retained and plotted.

### Machine Learning Model Construction

A binary classification model was created to distinguish Pre-RA participants from Non-converters at their baseline visit using both epigenetic markers and antibody levels. We first selected the feature set by taking the top 60 DMLs within each cell lineage based on p-value (empirical bayes-moderated t-test) between Pre-RA and Non-converter groups based on Visit 2 and At-Diagnosis visits, where available. A random seed was then set and Visit 1 samples were split into train and test subsets at a 70:30 ratio. The initial 60 DML feature set was ranked by importance based on the recursive feature elimination algorithm as implemented in the rfe() function in the R Caret^61^ package version 6.0.94. Only features with positive importance were retained.

Random Forest classification models were then trained on the training set with an increasing number of predictors, from most important to least, using 10-fold cross-validation and repeated 5 times. Each model was then tested on the unseen test set and evaluated based on accuracy. The train and test data split, feature importance evaluation, and model training and evaluation was repeated 100 times using different random seeds, from 1 to 100. The mean and standard deviation of the training and test accuracy was noted for each seed and each number of predictors.

We then repeated this process using only antibody/acute phase reactant data for each sample. Specifically, within this analysis anti-CCP3, C-Reactive Protein (CRP) and rheumatoid factors IgA, IgG, and IgM were included as features. Training and test prediction performance for DMLs in each individual cell lineage, as well as antibody/acute phase reactant only, were then plotted up to the predictor count that achieved the maximum mean training accuracy.

### Visualization

Principal components analysis (PCA) was used to visually portray the relationship between samples. This analysis was performed based on M-values using the prcomp() function in R. Hierarchical clustering and heatmaps were created using the R package pheatmap^62^ version 1.0.12 using M-values with outlier (top and bottom 1%) DMLs corrected. For longitudinal heatmaps depicting clusters of DMLs, DMLs were first clustered using hierarchical clustering. Clusters were then identified using the cutree() command from R package stats version 4.2.1. While individual probe methylation values were reported and visualized as β-values, all statistics were calculated using M-values. Additionally, all other visualization analyses were performed using M-values.

## Acknowledgements

The authors would like to express their gratitude to the study participants who made this study possible by contributing their biospecimens, data, and time. The authors also thank Dr. Jeffrey S. Carlin and the Benaroya Research Institute Clinical and Translational Cores for their support in study participant recruitment and evaluation.

## Author contributions

Conceptualization: GSF, KDD, VMH, WHR, JHB

Methodology: EBP, GSF, WW, DLB, KDD, WHR, JHB

Investigation: EBP, GSF, WW, DLB, KDD, VMH, WHR, JHB

Visualization: EBP

Funding acquisition: GSF, KDD, VMH, WHR, JHB

Project administration: GSF, KDD, VMH, WHR, JHB

Supervision: GSF, WW

Writing – original draft: EBP, GSF

Writing – review & editing: EBP, GSF, WW, DLB, KDD, VMH, WHR, JHB

## Data & Code Availability

The methylation data generated in this study have been deposited in NCBI’s Gene Expression Omnibus (accession number GSE277860). The code to reproduce all data analysis and create the figures in this study can be found at https://github.com/Wang-lab-UCSD/TIP-RA_longitudinal. Given the set of scripts and environments provided, only minimal changes (such as file paths) are necessary to reproduce the analyses of this work.

## References

(1) Holers, V. M.; Demoruelle, M. K.; Kuhn, K. A.; Buckner, J. H.; Robinson, W. H.; Okamoto, Y.; Norris, J. M.; Deane, K. D. Rheumatoid Arthritis and the Mucosal Origins Hypothesis: Protection Turns to Destruction. Nat. Rev. Rheumatol. 2018, 14 (9), 542–557. 10.1038/s41584-018-0070-0.

(2) Demoruelle, M. K.; Bowers, E.; Lahey, L. J.; Sokolove, J.; Purmalek, M.; Seto, N. L.; Weisman, M. H.; Norris, J. M.; Kaplan, M. J.; Holers, V. M.; Robinson, W. H.; Deane, K. D. Antibody Responses to Citrullinated and Non-Citrullinated Antigens in the Sputum of Subjects with and At-Risk for Rheumatoid Arthritis. Arthritis Rheumatol. Hoboken NJ 2018, 70 (4), 516–527. 10.1002/art.40401.

(3) Chriswell, M. E.; Lefferts, A. R.; Clay, M. R.; Hsu, A. R.; Seifert, J.; Feser, M. L.; Rims, C.; Bloom, M. S.; Bemis, E. A.; Liu, S.; Maerz, M. D.; Frank, D. N.; Demoruelle, M. K.; Deane, K. D.; James, E. A.; Buckner, J. H.; Robinson, W. H.; Holers, V. M.; Kuhn, K. A. Clonal IgA and IgG Autoantibodies from Individuals At-Risk for Rheumatoid Arthritis Identify an Arthritogenic Strain of Subdoligranulum. Sci. Transl. Med. 2022, 14 (668), eabn5166. 10.1126/scitranslmed.abn5166.

(4) Gravallese Ellen M.; Firestein Gary S. Rheumatoid Arthritis — Common Origins, Divergent Mechanisms. N. Engl. J. Med. 2023, 388 (6), 529–542. 10.1056/NEJMra2103726.

(5) Deane, K. D.; Holers, V. M. Rheumatoid Arthritis: Pathogenesis, Prediction and Prevention – An Emerging Paradigm Shift. Arthritis Rheumatol. Hoboken NJ 2021, 73 (2), 181–193. 10.1002/art.41417.

(6) James, E. A.; Holers, V. M.; Iyer, R.; Prideaux, E. B.; Rao, N. L.; Rims, C.; Muir, V. S.; Posso, S. E.; Bloom, M. S.; Zia, A.; Elliott, S. E.; Adamska, J. Z.; Ai, R.; Brewer, R. C.; Seifert, J. A.; Moss, L.; Barzideh, S.; Demoruelle, M. K.; Striebich, C. C.; Okamoto, Y.; Sainbayar, E.; Crook, A. A.; Peterson, R. A.; Vanderlinden, L. A.; Wang, W.; Boyle, D. L.; Robinson, W. H.; Buckner, J. H.; Firestein, G. S.; Deane, K. D. Multifaceted Immune Dysregulation Characterizes Individuals At-Risk for Rheumatoid Arthritis. Nat. Commun. 2023, 14 (1), 7637. 10.1038/s41467-023-43091-8.

(7) de la Calle-Fabregat, C.; Niemantsverdriet, E.; Cañete, J. D.; Li, T.; van der Helm-van Mil, H. M.; Rodríguez-Ubreva, J.; Ballestar, E. Prediction of the Progression of Undifferentiated Arthritis to Rheumatoid Arthritis Using DNA Methylation Profiling. Arthritis Rheumatol. 2021, 73 (12), 2229–2239. 10.1002/art.41885.

(8) Oytam, Y.; Sobhanmanesh, F.; Duesing, K.; Bowden, J. C.; Osmond-McLeod, M.; Ross, J. Risk-Conscious Correction of Batch Effects: Maximising Information Extraction from High-Throughput Genomic Datasets. BMC Bioinformatics 2016, 17 (1), 332. 10.1186/s12859-016-1212-5.

(9) Glossop, J. R.; Emes, R. D.; Nixon, N. B.; Packham, J. C.; Fryer, A. A.; Mattey, D. L.; Farrell, W. E. Genome-Wide Profiling in Treatment-Naive Early Rheumatoid Arthritis Reveals DNA Methylome Changes in T and B Lymphocytes. Epigenomics 2016, 8 (2), 209–224. 10.2217/epi.15.103.

(10) Nakazawa, M.; Ishii, H.; Aono, H.; Takai, M.; Honda, T.; Aratani, S.; Fukamizu, A.; Nakamura, H.; Yoshino, S.-I.; Kobata, T.; Nishioka, K.; Nakajima, T. Role of Notch-1 Intracellular Domain in Activation of Rheumatoid Synoviocytes. Arthritis Rheum. 2001, 44 (7), 1545–1554. 10.1002/1529-0131(200107)44:7<1545::AID-ART278>3.0.CO;2-Q.

(11) Kim, M. J.; Park, J.-S.; Lee, S. J.; Jang, J.; Park, J. S.; Back, S. H.; Bahn, G.; Park, J. H.; Kang, Y. M.; Kim, S. H.; Kwon, I. C.; Jo, D.-G.; Kim, K. Notch1 Targeting siRNA Delivery Nanoparticles for Rheumatoid Arthritis Therapy. J. Controlled Release 2015, 216, 140–148. 10.1016/j.jconrel.2015.08.025.

(12) Wei, K.; Korsunsky, I.; Marshall, J. L.; Gao, A.; Watts, G. F. M.; Major, T.; Croft, A. P.; Watts, J.; Blazar, P.; Lange, J.; Thornhill, T.; Filer, A.; Raza, K.; Donlin, L. T.; Consortium, A. M. P. R. arthritis and S. L. E. (AMP R.; Siebel, C. W.; Buckley, C. D.; Raychaudhuri, S.; Brenner, M. B. Notch Signaling Drives Synovial Fibroblast Identity and Arthritis Pathology. Nature 2020, 582 (7811), 259. 10.1038/s41586-020-2222-z.

(13) Simelyte, E.; Boyle, D.; Firestein, G. DNA Mismatch Repair Enzyme Expression in Synovial Tissue. Ann. Rheum. Dis. 2004, 63 (12), 1695. 10.1136/ard.2003.017210.

(14) Sendo, S.; Machado, C. R. L.; Boyle, D. L.; Benschop, R. J.; Perumal, N. B.; Choi, E.; Wang, W.; Firestein, G. S. Dysregulated and Neddylation Enhances Rheumatoid Arthritis Fibroblast-Like Synoviocyte Inflammatory Responses. Arthritis Rheumatol. 2024, 76 (8), 1252–1262. 10.1002/art.42856.

(15) Lassoued, S.; Moyano, C.; Beldjerd, M.; Pauly, P.; Lassoued, D.; Billey, T. Bortezomib Improved the Joint Manifestations of Rheumatoid Arthritis in Three Patients. Joint Bone Spine 2019, 86 (3), 381–382. 10.1016/j.jbspin.2019.01.019.

(16) Picard, R. R.; Berk, K. N. Data Splitting. Am. Stat. 1990, 44 (2), 140–147. 10.1080/00031305.1990.10475704.

(17) Ai, R.; Whitaker, J. W.; Boyle, D. L.; Tak, P. P.; Gerlag, D. M.; Wang, W.; Firestein, G. S. DNA Methylome Signature in Early Rheumatoid Arthritis Synoviocytes Compared with Longstanding Rheumatoid Arthritis Synoviocytes. Arthritis Rheumatol. Hoboken NJ 2015, 67 (7), 1978–1980. 10.1002/art.39123.

(18) Li Yim, A. Y. F.; Ferrero, E.; Maratou, K.; Lewis, H. D.; Royal, G.; Tough, D. F.; Larminie, C.; Mannens, M. M. A. M.; Henneman, P.; de Jonge, W. J.; van de Sande, M. G. H.; Gerlag, D. M.; Prinjha, R. K.; Tak, P. P. Novel Insights Into Rheumatoid Arthritis Through Characterization of Concordant Changes in DNA Methylation and Gene Expression in Synovial Biopsies of Patients With Differing Numbers of Swollen Joints. Front. Immunol. 2021, 12. 10.3389/fimmu.2021.651475.

(19) Pitaksalee, R.; Burska, A. N.; Ajaib, S.; Rogers, J.; Parmar, R.; Mydlova, K.; Xie, X.; Droop, A.; Nijjar, J. S.; Chambers, P.; Emery, P.; Hodgett, R.; McInnes, I. B.; Ponchel, F. Differential CpG DNA Methylation in Peripheral Naïve CD4+ T-Cells in Early Rheumatoid Arthritis Patients. Clin. Epigenetics 2020, 12, 54. 10.1186/s13148-020-00837-1.

(20) Karouzakis, E.; Raza, K.; Kolling, C.; Buckley, C. D.; Gay, S.; Filer, A.; Ospelt, C. Analysis of Early Changes in DNA Methylation in Synovial Fibroblasts of RA Patients before Diagnosis. Sci. Rep. 2018, 8, 7370. 10.1038/s41598-018-24240-2.

(21) Guderud, K.; Sunde, L. H.; Flåm, S. T.; Mæhlen, M. T.; Mjaavatten, M. D.; Lillegraven, S.; Aga, A.-B.; Evenrød, I. M.; Norli, E. S.; Andreassen, B. K.; Franzenburg, S.; Franke, A.; Haavardsholm, E. A.; Rayner, S.; Gervin, K.; Lie, B. A. Rheumatoid Arthritis Patients, Both Newly Diagnosed and Methotrexate Treated, Show More DNA Methylation Differences in CD4+ Memory Than in CD4+ Naïve T Cells. Front. Immunol. 2020, 11, 194. 10.3389/fimmu.2020.00194.

(22) Gosselt, H. R.; Vallerga, C. L.; Mandaviya, P. R.; Lubberts, E.; Hazes, J. M. W.; de Jonge, R.; Heil, S. G. Epigenome Wide Association Study of Response to Methotrexate in Early Rheumatoid Arthritis Patients. PLoS ONE 2021, 16 (3), e0247709. 10.1371/journal.pone.0247709.

(23) Rhead, B.; Holingue, C.; Cole, M.; Shao, X.; Quach, H. L.; Quach, D.; Shah, K.; Sinclair, E.; Graf, J.; Link, T.; Harrison, R.; Rahmani, E.; Halperin, E.; Wang, W.; Firestein, G. S.; Barcellos, L. F.; Criswell, L. A. Rheumatoid Arthritis Naive T Cells Share Hypermethylation Sites With Synoviocytes. Arthritis Rheumatol. Hoboken Nj 2017, 69 (3), 550–559. 10.1002/art.39952.

(24) Wang, K.; Liu, H.; Hu, Q.; Wang, L.; Liu, J.; Zheng, Z.; Zhang, W.; Ren, J.; Zhu, F.; Liu, G.-H. Epigenetic Regulation of Aging: Implications for Interventions of Aging and Diseases. Signal Transduct. Target. Ther. 2022, 7 (1), 1–22. 10.1038/s41392-022-01211-8.

(25) Grant, O. A.; Wang, Y.; Kumari, M.; Zabet, N. R.; Schalkwyk, L. Characterising Sex Differences of Autosomal DNA Methylation in Whole Blood Using the Illumina EPIC Array. Clin. Epigenetics 2022, 14 (1), 62. 10.1186/s13148-022-01279-7.

(26) Zeilinger, S.; Kühnel, B.; Klopp, N.; Baurecht, H.; Kleinschmidt, A.; Gieger, C.; Weidinger, S.; Lattka, E.; Adamski, J.; Peters, A.; Strauch, K.; Waldenberger, M.; Illig, T. Tobacco Smoking Leads to Extensive Genome-Wide Changes in DNA Methylation. PLoS ONE 2013, 8 (5), e63812. 10.1371/journal.pone.0063812.

(27) You, C.; Wu, S.; Zheng, S. C.; Zhu, T.; Jing, H.; Flagg, K.; Wang, G.; Jin, L.; Wang, S.; Teschendorff, A. E. A Cell-Type Deconvolution Meta-Analysis of Whole Blood EWAS Reveals Lineage-Specific Smoking-Associated DNA Methylation Changes. Nat. Commun. 2020, 11 (1), 4779. 10.1038/s41467-020-18618-y.

(28) Lyko, F. The DNA Methyltransferase Family: A Versatile Toolkit for Epigenetic Regulation. Nat. Rev. Genet. 2018, 19 (2), 81–92. 10.1038/nrg.2017.80.

(29) Li, M.; Wang, N.; Shen, Z.; Yan, J. Long Non-Coding RNA Growth Arrest-Specific Transcript 5 Regulates Rheumatoid Arthritis by Targeting Homeodomain-Interacting Protein Kinase 2. Clin. Exp. Rheumatol. 2020, 38 (6), 1145–1154.

(30) Cici, D.; Corrado, A.; Rotondo, C.; Cantatore, F. P. Wnt Signaling and Biological Therapy in Rheumatoid Arthritis and Spondyloarthritis. Int. J. Mol. Sci. 2019, 20 (22), 5552. 10.3390/ijms20225552.

(31) Carpenter, S.; Fitzgerald, K. A. Cytokines and Long Noncoding RNAs. Cold Spring Harb. Perspect. Biol. 2018, 10 (6), a028589. 10.1101/cshperspect.a028589.

(32) Paoletti, A.; Rohmer, J.; Ly, B.; Pascaud, J.; Rivière, E.; Seror, R.; Le Goff, B.; Nocturne, G.; Mariette, X. Monocyte/Macrophage Abnormalities Specific to Rheumatoid Arthritis Are Linked to miR-155 and Are Differentially Modulated by Different TNF Inhibitors. J. Immunol. 2019, 203 (7), 1766–1775. 10.4049/jimmunol.1900386.

(33) Yan, S.; Wang, P.; Wang, J.; Yang, J.; Lu, H.; Jin, C.; Cheng, M.; Xu, D. Long Non-Coding RNA HIX003209 Promotes Inflammation by Sponging miR-6089 via TLR4/NF-κB Signaling Pathway in Rheumatoid Arthritis. Front. Immunol. 2019, 10. 10.3389/fimmu.2019.02218.

(34) Rodríguez-Ubreva, J.; De La Calle-Fabregat, C.; Li, T.; Ciudad, L.; Ballestar, M. L.; Català-Moll, F.; Morante-Palacios, O.; Garcia-Gomez, A.; Celis, R.; Humby, F.; Nerviani, A.; Martin, J.; Pitzalis, C.; Cañete, J. D.; Ballestar, E. Inflammatory Cytokines Shape a Changing DNA Methylome in Monocytes Mirroring Disease Activity in Rheumatoid Arthritis. Ann. Rheum. Dis. 2019, 78 (11), 1505–1516. 10.1136/annrheumdis-2019-215355.

(35) Pidsley, R.; Zotenko, E.; Peters, T. J.; Lawrence, M. G.; Risbridger, G. P.; Molloy, P.; Van Djik, S.; Muhlhausler, B.; Stirzaker, C.; Clark, S. J. Critical Evaluation of the Illumina MethylationEPIC BeadChip Microarray for Whole-Genome DNA Methylation Profiling. Genome Biol. 2016, 17, 208. 10.1186/s13059-016-1066-1.

(36) Ferreté-Bonastre, A. G.; Cortés-Hernández, J.; Ballestar, E. What Can We Learn from DNA Methylation Studies in Lupus? Clin. Immunol. 2022, 234, 108920. 10.1016/j.clim.2021.108920.

(37) Ross, M. T.; Grafham, D. V.; Coffey, A. J.; Scherer, S.; McLay, K.; Muzny, D.; Platzer, M.; Howell, G. R.; Burrows, C.; Bird, C. P.; Frankish, A.; Lovell, F. L.; Howe, K. L.; Ashurst, J. L.; Fulton, R. S.; Sudbrak, R.; Wen, G.; Jones, M. C.; Hurles, M. E.; Andrews, T. D.; Scott, C. E.; Searle, S.; Ramser, J.; Whittaker, A.; Deadman, R.; Carter, N. P.; Hunt, S. E.; Chen, R.; Cree, A.; Gunaratne, P.; Havlak, P.; Hodgson, A.; Metzker, M. L.; Richards, S.; Scott, G.; Steffen, D.; Sodergren, E.; Wheeler, D. A.; Worley, K. C.; Ainscough, R.; Ambrose, K. D.; Ansari-Lari, M. A.; Aradhya, S.; Ashwell, R. I. S.; Babbage, A. K.; Bagguley, C. L.; Ballabio, A.; Banerjee, R.; Barker, G. E.; Barlow, K. F.; Barrett, I. P.; Bates, K. N.; Beare, D. M.; Beasley, H.; Beasley, O.; Beck, A.; Bethel, G.; Blechschmidt, K.; Brady, N.; Bray-Allen, S.; Bridgeman, A. M.; Brown, A. J.; Brown, M. J.; Bonnin, D.; Bruford, E. A.; Buhay, C.; Burch, P.; Burford, D.; Burgess, J.; Burrill, W.; Burton, J.; Bye, J. M.; Carder, C.; Carrel, L.; Chako, J.; Chapman, J. C.; Chavez, D.; Chen, E.; Chen, G.; Chen, Y.; Chen, Z.; Chinault, C.; Ciccodicola, A.; Clark, S. Y.; Clarke, G.; Clee, C. M.; Clegg, S.; Clerc-Blankenburg, K.; Clifford, K.; Cobley, V.; Cole, C. G.; Conquer, J. S.; Corby, N.; Connor, R. E.; David, R.; Davies, J.; Davis, C.; Davis, J.; Delgado, O.; DeShazo, D.; Dhami, P.; Ding, Y.; Dinh, H.; Dodsworth, S.; Draper, H.; Dugan-Rocha, S.; Dunham, A.; Dunn, M.; Durbin, K. J.; Dutta, I.; Eades, T.; Ellwood, M.; Emery-Cohen, A.; Errington, H.; Evans, K. L.; Faulkner, L.; Francis, F.; Frankland, J.; Fraser, A. E.; Galgoczy, P.; Gilbert, J.; Gill, R.; Glöckner, G.; Gregory, S. G.; Gribble, S.; Griffiths, C.; Grocock, R.; Gu, Y.; Gwilliam, R.; Hamilton, C.; Hart, E. A.; Hawes, A.; Heath, P. D.; Heitmann, K.; Hennig, S.; Hernandez, J.; Hinzmann, B.; Ho, S.; Hoffs, M.; Howden, P. J.; Huckle, E. J.; Hume, J.; Hunt, P. J.; Hunt, A. R.; Isherwood, J.; Jacob, L.; Johnson, D.; Jones, S.; de Jong, P. J.; Joseph, S. S.; Keenan, S.; Kelly, S.; Kershaw, J. K.; Khan, Z.; Kioschis, P.; Klages, S.; Knights, A. J.; Kosiura, A.; Kovar-Smith, C.; Laird, G. K.; Langford, C.; Lawlor, S.; Leversha, M.; Lewis, L.; Liu, W.; Lloyd, C.; Lloyd, D. M.; Loulseged, H.; Loveland, J. E.; Lovell, J. D.; Lozado, R.; Lu, J.; Lyne, R.; Ma, J.; Maheshwari, M.; Matthews, L. H.; McDowall, J.; McLaren, S.; McMurray, A.; Meidl, P.; Meitinger, T.; Milne, S.; Miner, G.; Mistry, S. L.; Morgan, M.; Morris, S.; Müller, I.; Mullikin, J. C.; Nguyen, N.; Nordsiek, G.; Nyakatura, G.; O’Dell, C. N.; Okwuonu, G.; Palmer, S.; Pandian, R.; Parker, D.; Parrish, J.; Pasternak, S.; Patel, D.; Pearce, A. V.; Pearson, D. M.; Pelan, S. E.; Perez, L.; Porter, K. M.; Ramsey, Y.; Reichwald, K.; Rhodes, S.; Ridler, K. A.; Schlessinger, D.; Schueler, M. G.; Sehra, H. K.; Shaw-Smith, C.; Shen, H.; Sheridan, E. M.; Shownkeen, R.; Skuce, C. D.; Smith, M. L.; Sotheran, E. C.; Steingruber, H. E.; Steward, C. A.; Storey, R.; Swann, R. M.; Swarbreck, D.; Tabor, P. E.; Taudien, S.; Taylor, T.; Teague, B.; Thomas, K.; Thorpe, A.; Timms, K.; Tracey, A.; Trevanion, S.; Tromans, A. C.; d’Urso, M.; Verduzco, D.; Villasana, D.; Waldron, L.; Wall, M.; Wang, Q.; Warren, J.; Warry, G. L.; Wei, X.; West, A.; Whitehead, S. L.; Whiteley, M. N.; Wilkinson, J. E.; Willey, D. L.; Williams, G.; Williams, L.; Williamson, A.; Williamson, H.; Wilming, L.; Woodmansey, R. L.; Wray, P. W.; Yen, J.; Zhang, J.; Zhou, J.; Zoghbi, H.; Zorilla, S.; Buck, D.; Reinhardt, R.; Poustka, A.; Rosenthal, A.; Lehrach, H.; Meindl, A.; Minx, P. J.; Hillier, L. W.; Willard, H. F.; Wilson, R. K.; Waterston, R. H.; Rice, C. M.; Vaudin, M.; Coulson, A.; Nelson, D. L.; Weinstock, G.; Sulston, J. E.; Durbin, R.; Hubbard, T.; Gibbs, R. A.; Beck, S.; Rogers, J.; Bentley, D. R. The DNA Sequence of the Human X Chromosome. Nature 2005, 434 (7031), 325–337. 10.1038/nature03440.

(38) McComish, J.; Mundy, J.; Sullivan, T.; Proudman, S. M.; Hissaria, P. Changes in Peripheral Blood B Cell Subsets at Diagnosis and after Treatment with Disease-Modifying Anti-Rheumatic Drugs in Patients with Rheumatoid Arthritis: Correlation with Clinical and Laboratory Parameters. Int. J. Rheum. Dis. 2015, 18 (4), 421–432. 10.1111/1756-185X.12325.

(39) Wang, Y.; Lloyd, K. A.; Melas, I.; Zhou, D.; Thyagarajan, R.; Lindqvist, J.; Hansson, M.; Svärd, A.; Mathsson-Alm, L.; Kastbom, A.; Lundberg, K.; Klareskog, L.; Catrina, A. I.; Rapecki, S.; Malmström, V.; Grönwall, C. Rheumatoid Arthritis Patients Display B-Cell Dysregulation Already in the Naïve Repertoire Consistent with Defects in B-Cell Tolerance. Sci. Rep. 2019, 9 (1), 19995. 10.1038/s41598-019-56279-0.

(40) Zhou, B.; Lin, W.; Long, Y.; Yang, Y.; Zhang, H.; Wu, K.; Chu, Q. Notch Signaling Pathway: Architecture, Disease, and Therapeutics. Signal Transduct. Target. Ther. 2022, 7 (1), 1–33. 10.1038/s41392-022-00934-y.

(41) Ferro, E. S.; Gewehr, M. C. F.; Navon, A. Thimet Oligopeptidase Biochemical and Biological Significances: Past, Present, and Future Directions. Biomolecules 2020, 10 (9), 1229. 10.3390/biom10091229.

(42) Kim, S. I.; Pabon, A.; Swanson, T. A.; Glucksman, M. J. Regulation of Cell-Surface Major Histocompatibility Complex Class I Expression by the Endopeptidase EC3.4.24.15 (Thimet Oligopeptidase). Biochem. J. 2003, 375 (Pt 1), 111–120. 10.1042/BJ20030490.

(43) York, I. A.; Mo, A. X. Y.; Lemerise, K.; Zeng, W.; Shen, Y.; Abraham, C. R.; Saric, T.; Goldberg, A. L.; Rock, K. L. The Cytosolic Endopeptidase, Thimet Oligopeptidase, Destroys Antigenic Peptides and Limits the Extent of MHC Class I Antigen Presentation. Immunity 2003, 18 (3), 429–440. 10.1016/s1074-7613(03)00058-x.

(44) Guinan, A. F.; Rochfort, K. D.; Fitzpatrick, P. A.; Walsh, T. G.; Pierotti, A. R.; Phelan, S.; Murphy, R. P.; Cummins, P. M. Shear Stress Is a Positive Regulator of Thimet Oligopeptidase (EC3.4.24.15) in Vascular Endothelial Cells: Consequences for MHC1 Levels. Cardiovasc. Res. 2013, 99 (3), 545–554. 10.1093/cvr/cvt127.

(45) Shchetynsky, K.; Diaz-Gallo, L.-M.; Folkersen, L.; Hensvold, A. H.; Catrina, A. I.; Berg, L.; Klareskog, L.; Padyukov, L. Discovery of New Candidate Genes for Rheumatoid Arthritis through Integration of Genetic Association Data with Expression Pathway Analysis. Arthritis Res. Ther. 2017, 19, 19. 10.1186/s13075-017-1220-5.

(46) Aryee, M. J.; Jaffe, A. E.; Corrada-Bravo, H.; Ladd-Acosta, C.; Feinberg, A. P.; Hansen, K. D.; Irizarry, R. A. Minfi: A Flexible and Comprehensive Bioconductor Package for the Analysis of Infinium DNA Methylation Microarrays. Bioinformatics 2014, 30 (10), 1363–1369. 10.1093/bioinformatics/btu049.

(47) Inkster, A. M.; Wong, M. T.; Matthews, A. M.; Brown, C. J.; Robinson, W. P. Who’s Afraid of the X? Incorporating the X and Y Chromosomes into the Analysis of DNA Methylation Array Data. Epigenetics Chromatin 2023, 16 (1), 1. 10.1186/s13072-022-00477-0.

(48) Daca-Roszak, P.; Pfeifer, A.; Żebracka-Gala, J.; Rusinek, D.; Szybińska, A.; Jarząb, B.; Witt, M.; Ziętkiewicz, E. Impact of SNPs on Methylation Readouts by Illumina Infinium HumanMethylation450 BeadChip Array: Implications for Comparative Population Studies. BMC Genomics 2015, 16, 1003. 10.1186/s12864-015-2202-0.

(49) Chen, Y.; Lemire, M.; Choufani, S.; Butcher, D. T.; Grafodatskaya, D.; Zanke, B. W.; Gallinger, S.; Hudson, T. J.; Weksberg, R. Discovery of Cross-Reactive Probes and Polymorphic CpGs in the Illumina Infinium HumanMethylation450 Microarray. Epigenetics 2013, 8 (2), 203–209. 10.4161/epi.23470.

(50) IlluminaHumanMethylationEPICanno.ilm10b4.hg19. Bioconductor. http://bioconductor.org/packages/IlluminaHumanMethylationEPICanno.ilm10b4.hg19/ (accessed 2024-05-06).

(51) LaBarre, B. A.; Goncearenco, A.; Petrykowska, H. M.; Jaratlerdsiri, W.; Bornman, M. S. R.; Hayes, V. M.; Elnitski, L. MethylToSNP: Identifying SNPs in Illumina DNA Methylation Array Data. Epigenetics Chromatin 2019, 12 (1), 79. 10.1186/s13072-019-0321-6.

(52) Du, P.; Zhang, X.; Huang, C.-C.; Jafari, N.; Kibbe, W. A.; Hou, L.; Lin, S. M. Comparison of Beta-Value and M-Value Methods for Quantifying Methylation Levels by Microarray Analysis. BMC Bioinformatics 2010, 11 (1), 587. 10.1186/1471-2105-11-587.

(53) Ritchie, M. E.; Phipson, B.; Wu, D.; Hu, Y.; Law, C. W.; Shi, W.; Smyth, G. K. Limma Powers Differential Expression Analyses for RNA-Sequencing and Microarray Studies. Nucleic Acids Res. 2015, 43 (7), e47. 10.1093/nar/gkv007.

(54) Church, D. M.; Schneider, V. A.; Graves, T.; Auger, K.; Cunningham, F.; Bouk, N.; Chen, H.-C.; Agarwala, R.; McLaren, W. M.; Ritchie, G. R. S.; Albracht, D.; Kremitzki, M.; Rock, S.; Kotkiewicz, H.; Kremitzki, C.; Wollam, A.; Trani, L.; Fulton, L.; Fulton, R.; Matthews, L.; Whitehead, S.; Chow, W.; Torrance, J.; Dunn, M.; Harden, G.; Threadgold, G.; Wood, J.; Collins, J.; Heath, P.; Griffiths, G.; Pelan, S.; Grafham, D.; E. Eichler, E.; Weinstock, G.; Mardis, E. R.; Wilson, R. K.; Howe, K.; Flicek, P.; Hubbard, T. Modernizing Reference Genome Assemblies. PLoS Biol. 2011, 9 (7), e1001091. 10.1371/journal.pbio.1001091.

(55) Durinck, S.; Spellman, P. T.; Birney, E.; Huber, W. Mapping Identifiers for the Integration of Genomic Datasets with the R/Bioconductor Package biomaRt. Nat. Protoc. 2009, 4 (8), 1184–1191. 10.1038/nprot.2009.97.

(56) Yu, G.; He, Q.-Y. ReactomePA: An R/Bioconductor Package for Reactome Pathway Analysis and Visualization. Mol. Biosyst. 2016, 12 (2), 477–479. 10.1039/c5mb00663e.

(57) Croft, D.; O’Kelly, G.; Wu, G.; Haw, R.; Gillespie, M.; Matthews, L.; Caudy, M.; Garapati, P.; Gopinath, G.; Jassal, B.; Jupe, S.; Kalatskaya, I.; Mahajan, S.; May, B.; Ndegwa, N.; Schmidt, E.; Shamovsky, V.; Yung, C.; Birney, E.; Hermjakob, H.; D’Eustachio, P.; Stein, L. Reactome: A Database of Reactions, Pathways and Biological Processes. Nucleic Acids Res. 2011, 39 (Database issue), D691–D697. 10.1093/nar/gkq1018.

(58) Fabregat, A.; Sidiropoulos, K.; Viteri, G.; Forner, O.; Marin-Garcia, P.; Arnau, V.; D’Eustachio, P.; Stein, L.; Hermjakob, H. Reactome Pathway Analysis: A High-Performance in-Memory Approach. BMC Bioinformatics 2017, 18 (1), 142. 10.1186/s12859-017-1559-2.

(59) Bernstein, B. E.; Stamatoyannopoulos, J. A.; Costello, J. F.; Ren, B.; Milosavljevic, A.; Meissner, A.; Kellis, M.; Marra, M. A.; Beaudet, A. L.; Ecker, J. R.; Farnham, P. J.; Hirst, M.; Lander, E. S.; Mikkelsen, T. S.; Thomson, J. A. The NIH Roadmap Epigenomics Mapping Consortium. Nat. Biotechnol. 2010, 28 (10), 1045–1048. 10.1038/nbt1010-1045.

(60) Ernst, J.; Kellis, M. ChromHMM: Automating Chromatin-State Discovery and Characterization. Nat. Methods 2012, 9 (3), 215–216. 10.1038/nmeth.1906.

(61) Kuhn, M. Building Predictive Models in R Using the Caret Package. J. Stat. Softw. 2008, 28, 1–26. 10.18637/jss.v028.i05.

(62) Kolde, R. Pheatmap: Pretty Heatmaps, 2019. https://cran.r-project.org/web/packages/pheatmap/index.html (accessed 2024-05-06).

