## Supplemental Materials for "Epigenetic trajectory predicts development of clinical rheumatoid arthritis in anti-citrullinated protein antibody positive individuals: Targeting Immune Responses for Prevention of Rheumatoid Arthritis (TIP-RA)"

**Supplementary Figure 1.** Overview of TIP-RA study design and data availability.

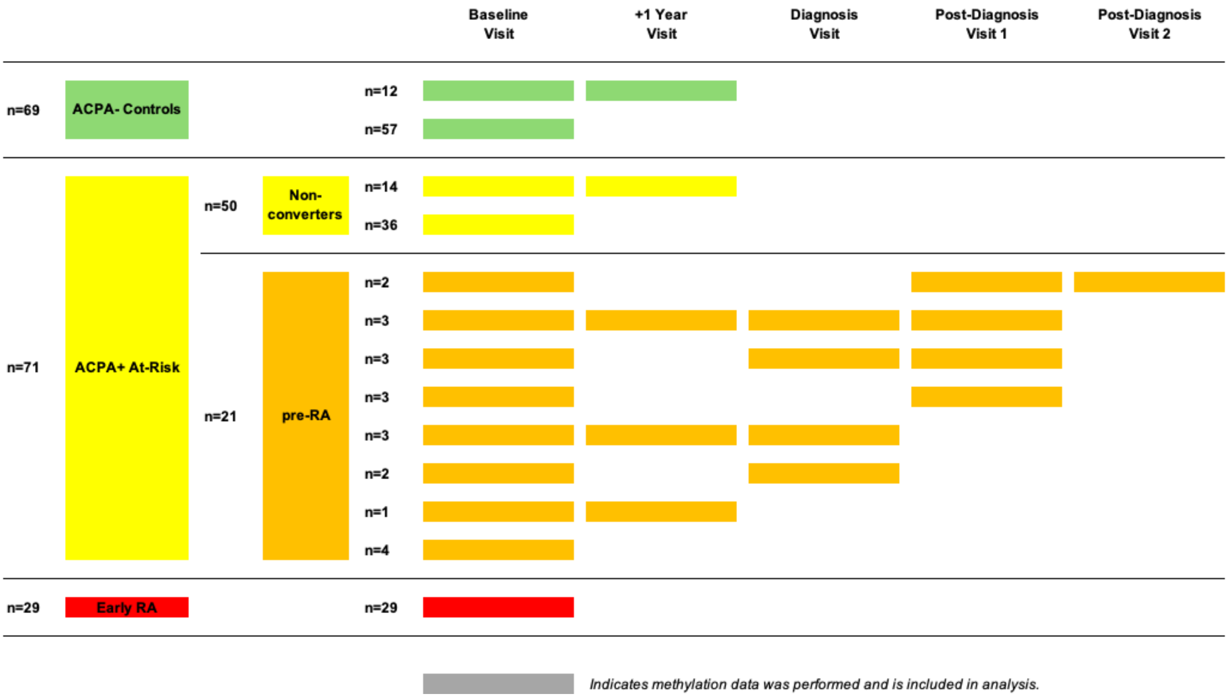

**Supplementary Figure 2.** PCA of filtered autosomal loci before batch correction.

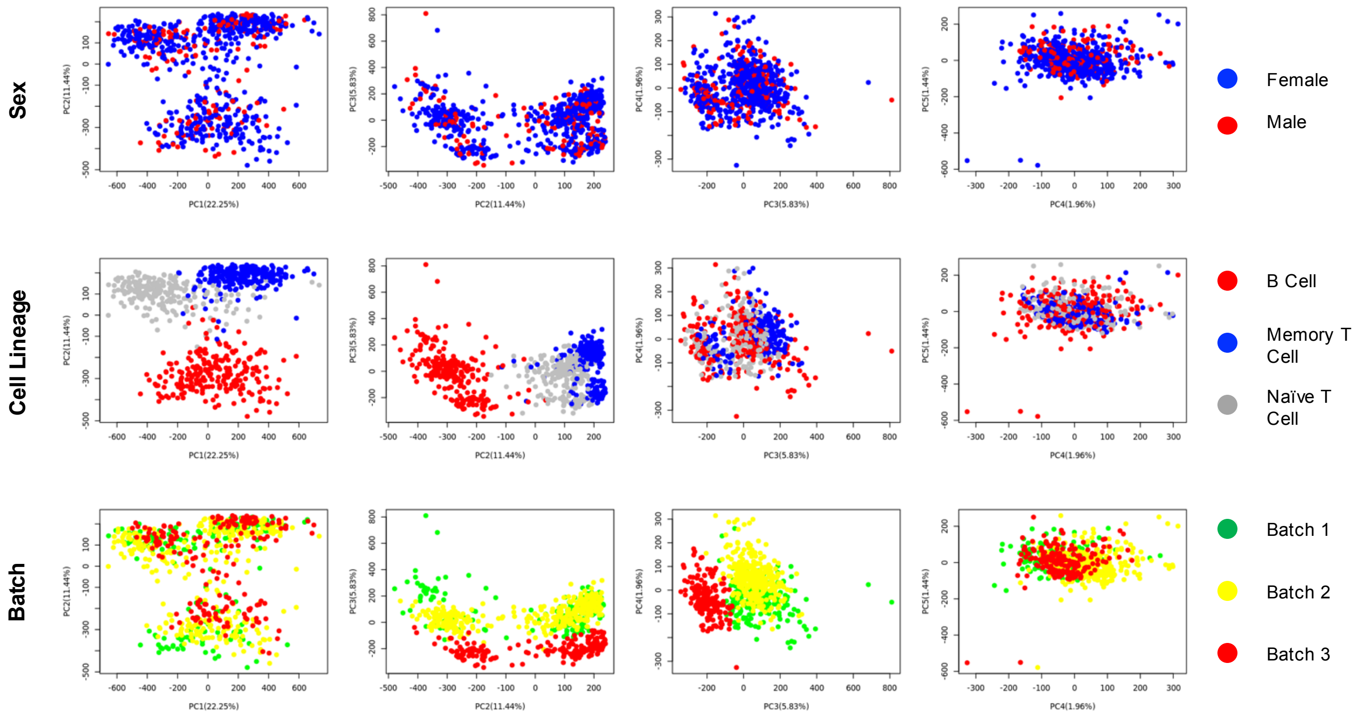

**Supplementary Figure 3.** PCA of filtered autosomal loci after batch correction.

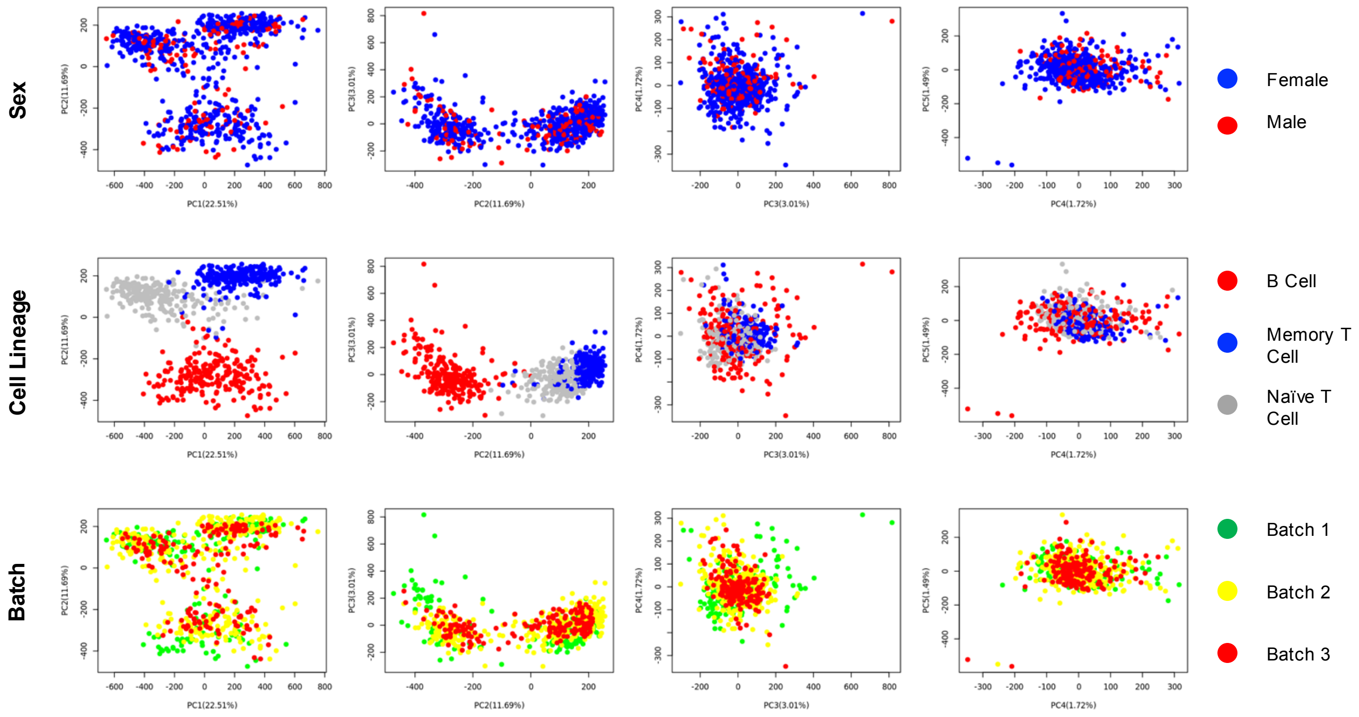

**Supplementary Figure 4.** Heatmap of DMLs identified in paired analysis of pre-diagnosis to diagnosis samples, diagnosis to post-diagnosis samples and pre-diagnosis to post-diagnosis samples for CCP3+ Converter patients.

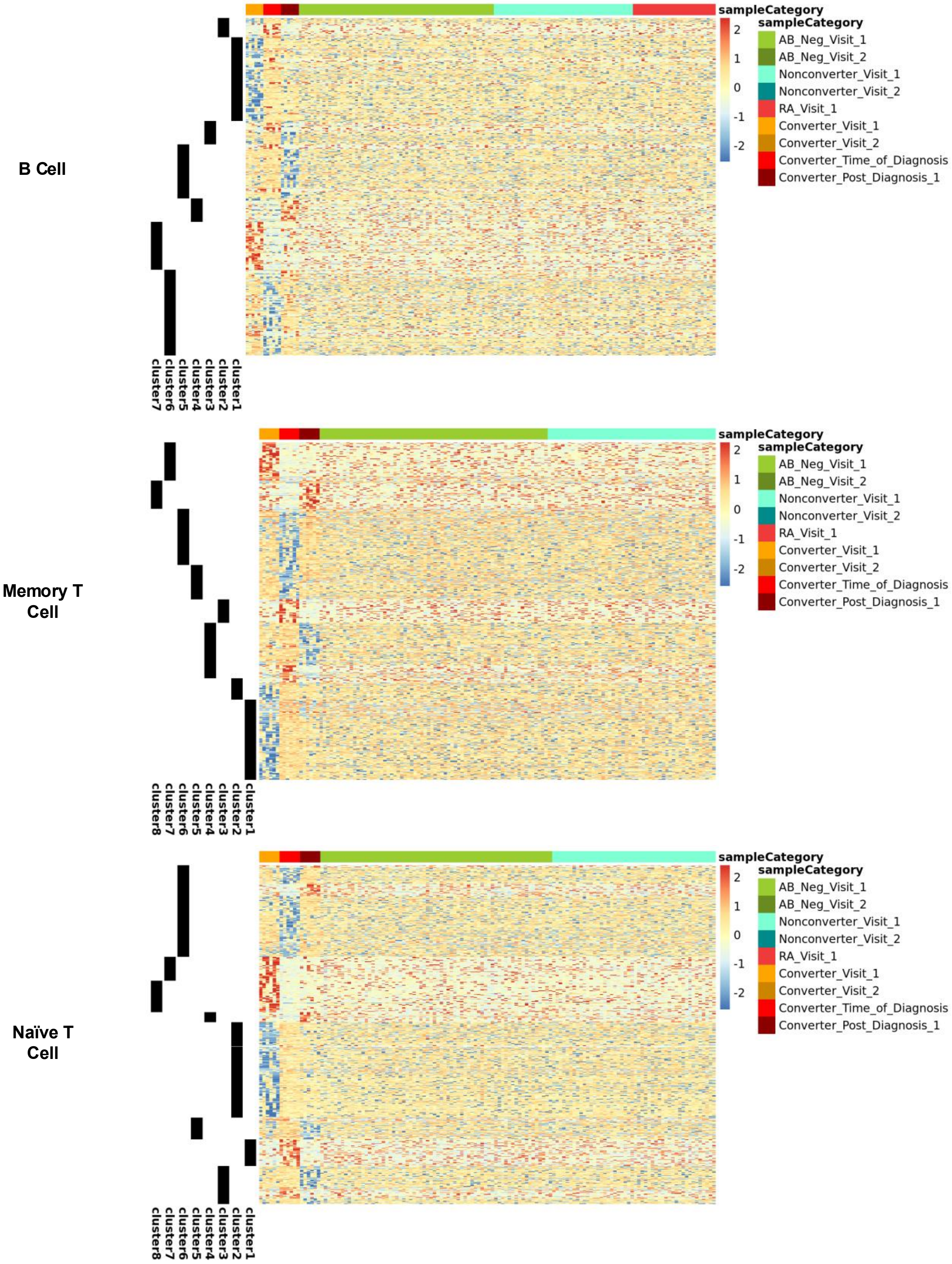

**Supplementary Figure 5.** Prediction accuracy of Random Forest models based on DML methylation levels from each cell lineage and antibody levels.

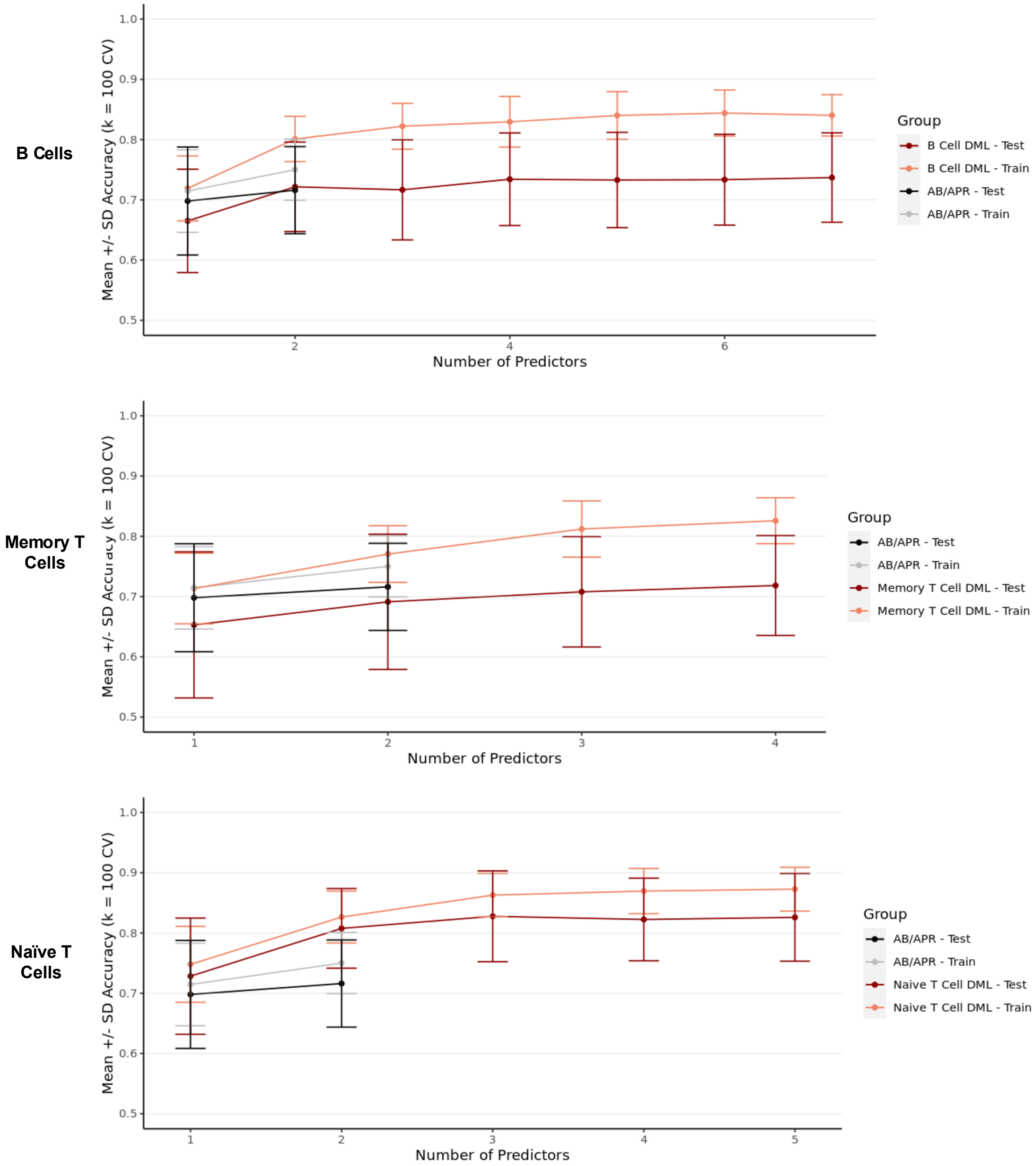

**Supplementary Figure 6.** Beta values of probes used in Random Forest predictions for each cell type. No probes have patterns suggesting SNP overlap.

**B Cell Predictors**

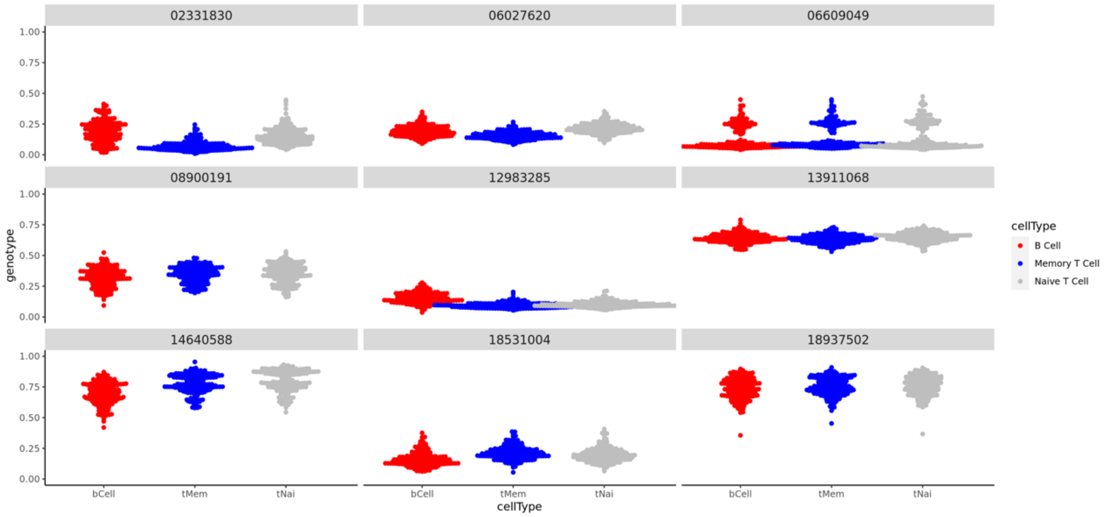


**Memory T Cell Predictors**

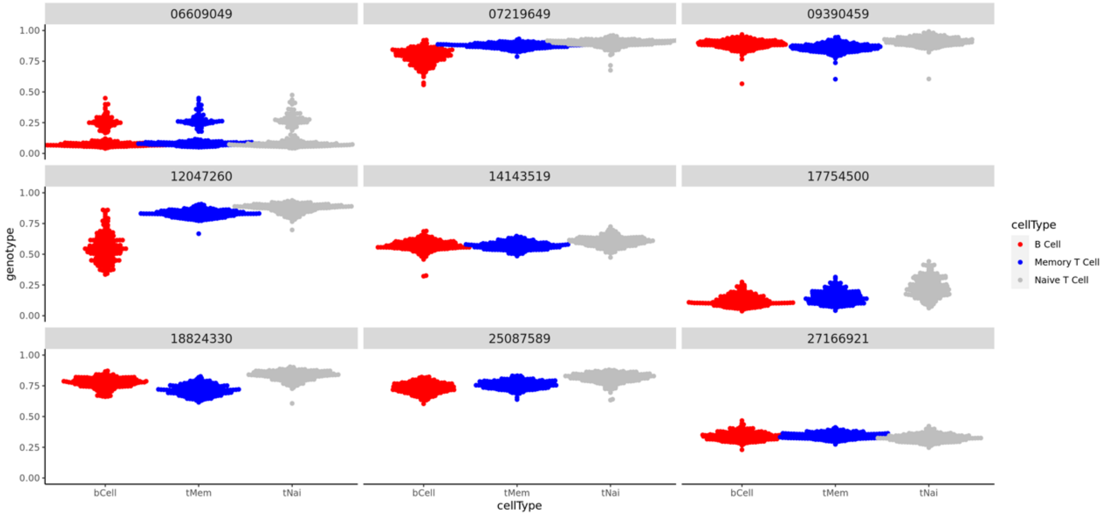


**Naïve T Cell Predictors**

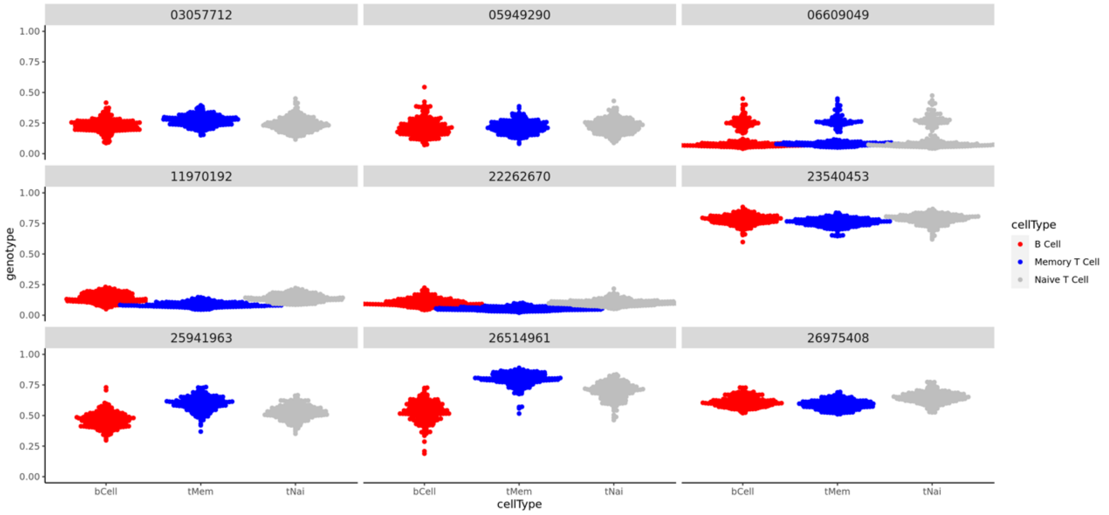

**Supplementary Figure 7.** Methylation intensities (depicted as beta values) of the three most important DML predictors models derived from naïve T cells samples. All points shown are derived from naïve T cell samples. Each color indicates a specific patient.

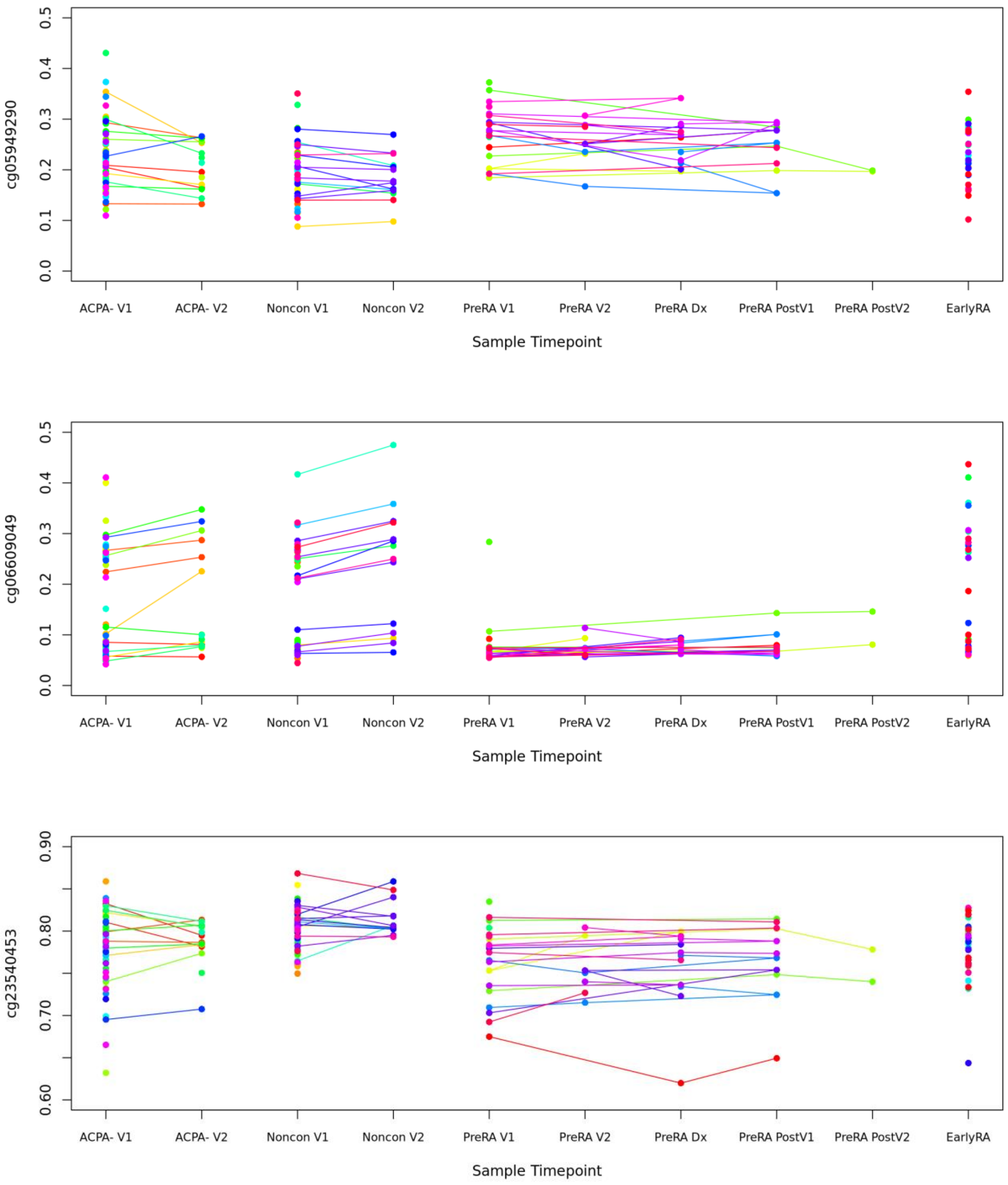

**Supplementary Table 1.** The number of DMLs / DMGs identified in cross sectional analysis among CCP3- controls, CCP3+ Nonconverter, CCP3+ Pre-RA, and Early RA in B cell, memory T cell, naïve T cell, and pooled lineage (in DMLs/DMGs), based on P-Value < 0.05, Difference of M > 0.5.

| | CCP3- vs Noncon | CCP3- vs Pre-RA | CCP3- vs RA | Noncon vs Pre-RA | Noncon vs RA | Pre-RA vs RA |
|---|---|---|---|---|---|---|
| **B Cell** | 3 / 1 | 166 / 91 | 93 / 42 | 202 / 92 | 153 / 63 | 360 / 180 |
| **Memory T Cell** | 3 / 2 | 66 / 23 | 54 / 27 | 89 / 36 | 86 / 35 | 276 / 147 |
| **Naive T Cell** | 4 / 2 | 83 / 35 | 61 / 21 | 37 / 71 | 100 / 41 | 292 / 154 |
| **Pooled Lineage** | 0 / 0 | 5 / 6 | 0 / 0 | 0 / 0 | 3 / 1 | 7 / 12 |

**Supplementary Table 2.** Pathway enrichment tables of baseline cross-sectional analysis in B cell samples. DMGs mapped from DMLs identified in comparison of CCP3+ Nonconverters and CCP3+ Pre-RA.

| Pathway | FDR | Gene Symbol |
|---|---|---|
| NOTCH1 Intracellular Domain Regulates Transcription | 4.94E-02 | HDAC2/MYC/HES1 |
| Nucleotide Excision Repair | 4.94E-02 | AQR/ERCC1/RUVBL1/RFC1 |
| Inositol phosphate metabolism | 4.94E-02 | INPP5A/PLCD1/NUDT3 |
| Suppression of phagosomal maturation | 4.94E-02 | KPNA1/RAB5A |
| Signaling by NOTCH1 PEST Domain Mutants in Cancer | 4.94E-02 | HDAC2/MYC/HES1 |
| Signaling by NOTCH1 in Cancer | 4.94E-02 | HDAC2/MYC/HES1 |
| Constitutive Signaling by NOTCH1 PEST Domain Mutants | 4.94E-02 | HDAC2/MYC/HES1 |
| Signaling by NOTCH1 HD+PEST Domain Mutants in Cancer | 4.94E-02 | HDAC2/MYC/HES1 |
| Constitutive Signaling by NOTCH1 HD+PEST Domain Mutants | 4.94E-02 | HDAC2/MYC/HES1 |
| Dual incision in TC-NER | 6.46E-02 | AQR/ERCC1/RFC1 |
| Signaling by NOTCH1 | 8.14E-02 | HDAC2/MYC/HES1 |
| Transcription-Coupled Nucleotide Excision Repair (TC-NER) | 8.42E-02 | AQR/ERCC1/RFC1 |
| Response of Mtb to phagocytosis | 8.42E-02 | KPNA1/RAB5A |
| Global Genome Nucleotide Excision Repair (GG-NER) | 9.15E-02 | ERCC1/RUVBL1/RFC1 |
| Signaling by ALK | 9.15E-02 | HDAC2/MYC |
| Infection with Mycobacterium tuberculosis | 9.15E-02 | KPNA1/RAB5A |
| RAB GEFs exchange GTP for GDP on RABs | 9.15E-02 | RAB3GAP1/RAB5A/AKT3 |

**Supplementary Table 3.** Pathway enrichment tables of paired longitudinal analysis of B cell samples.

### Cluster 1

| Pathway | -log(FDR) | Gene Symbol |
|---|---|---|
| UCH proteinases | 1.72 | ACTL6A/PSMD14/SENP8/PSMB9 |
| Cell Cycle Checkpoints | 1.32 | GTSE1/PSMD14/CENPL/PSMB9 |
| Deubiquitination | 1.32 | ACTL6A/PSMD14/SENP8/PSMB9 |
| The role of GTSE1 in G2/M progression after G2 checkpoint | 1.32 | GTSE1/PSMD14/PSMB9 |
| Degradation of beta-catenin by the destruction complex | 1.32 | PSMD14/TLE4/PSMB9 |
| ER-Phagosome pathway | 1.32 | PSMD14/PSMB9/VAMP3 |
| Antigen processing-Cross presentation | 1.32 | PSMD14/PSMB9/VAMP3 |
| PTEN Regulation | 1.32 | ATN1/PSMD14/PSMB9 |
| G2/M Checkpoints | 1.32 | GTSE1/PSMD14/PSMB9 |
| Separation of Sister Chromatids | 1.32 | PSMD14/CENPL/PSMB9 |

### Cluster 2

| Pathway | -log(FDR) | Gene Symbol |
|---|---|---|
| KSRP (KHSRP) binds and destabilizes mRNA | 1.05 | EXOSC7/MAPK11 |
| ERK/MAPK targets | 1.05 | PPP2CA/MAPK11 |

### Cluster 4

| Pathway | -log(FDR) | Gene Symbol |
|---|---|---|
| Mitotic Prometaphase | 1.03 | NCAPD2/NEDD1 |

### Cluster 7

| Pathway | -log(FDR) | Gene Symbol |
|---|---|---|
| Deadenylation-dependent mRNA decay | 1.16 | EXOSC1/EIF4E |
| Anchoring of the basal body to the plasma membrane | 1.00 | TUBG1/NPHP4 |
| Cilium Assembly | 1.00 | TUBG1/NPHP4 |

**Supplementary Table 4.** Pathway enrichment tables of paired longitudinal analysis of memory T cell samples.

### Cluster 7

| Pathway | -log(FDR) | Gene Symbol |
|---|---|---|
| Neurotransmitter receptors and postsynaptic signal transmission | 1.17 | CAMK2G/GABBR1 |
| Transmission across Chemical Synapses | 1.17 | CAMK2G/GABBR1 |
| G alpha (i) signalling events | 1.17 | CAMK2G/GABBR1 |
| RHO GTPase Effectors | 1.17 | DVL3/KTN1 |
| Neuronal System | 1.17 | CAMK2G/GABBR1 |

**Supplementary Table 5.** Pathway enrichment tables of paired longitudinal analysis of naïve T cell samples.

#### All DMGs

| Pathway | -log(FDR) | Gene Symbol |
|---|---|---|
| Caspase activation via extrinsic apoptotic signalling pathway | 1.09 | RIPK1/TICAM2/TMED7-TICAM2/UNC5A |
| TRIF-mediated programmed cell death | 1.09 | RIPK1/TICAM2/TMED7-TICAM2 |

#### Cluster 1

| Pathway | -log(FDR) | Gene Symbol |
|---|---|---|
| SLC-mediated transmembrane transport | 1.11 | SLC28A3/SLC12A4 |

#### Cluster 2

| Pathway | -log(FDR) | Gene Symbol |
|---|---|---|
| Ion channel transport | 1.87 | CUTC/FKBP1B/ATP11B/CALM2/TRPC1/C8orf44-SGK3 |
| Stimuli-sensing channels | 1.14 | FKBP1B/CALM2/TRPC1/C8orf44-SGK3 |
| Ion homeostasis | 1.14 | FKBP1B/CALM2/TRPC1 |
| Ion transport by P-type ATPases | 1.14 | CUTC/ATP11B/CALM2 |
| Glycogen breakdown (glycogenolysis) | 1.12 | CALM2/PYGL |
| Protein methylation | 1.05 | EEF1A1/CALM2 |

#### Cluster 7

| Pathway | -log(FDR) | Gene Symbol |
|---|---|---|
| Signaling by Nuclear Receptors | 1.25 | PRKCZ/ANGPTL3/ESR2 |
| Extra-nuclear estrogen signaling | 1.25 | PRKCZ/ESR2 |
| Visual phototransduction | 1.18 | AGRN/CNGA1 |
| Signaling by TGFB family members | 1.18 | PRKCZ/BMPR1A |
| ESR-mediated signaling | 1.18 | PRKCZ/ESR2 |
| Diseases of metabolism | 1.18 | AGRN/OPLAH |

#### Cluster 8

| Pathway | -log(FDR) | Gene Symbol |
|---|---|---|
| G2/M Transition | 1.08 | TUBGCP5/PPP1R12A |
| Mitotic G2-G2/M phases | 1.08 | TUBGCP5/PPP1R12A |
